# Neurons enhance blood-brain barrier function via upregulating claudin-5 and VE-cadherin expression due to GDNF secretion

**DOI:** 10.1101/2024.02.07.579396

**Authors:** Lu Yang, Zijin Lin, Ruijing Mu, Wenhan Wu, Hao Zhi, Xiaodong Liu, Hanyu Yang, Li Liu

**Affiliations:** Department of Pharmacology, School of Pharmacy, China Pharmaceutical University, Nanjing 210009, China

**Keywords:** Blood-Brain Barrier, *in vitro* BBB model, Glia-derived neurotrophic factor, IVIVC, Co-culture, Tight junction

## Abstract

Blood-brain barrier (BBB) prevents neurotoxins from entering central nervous system. We aimed to establish and characterize an *in vitro* triple co-culture BBB model consisting of brain endothelial cells hCMEC/D3, astrocytoma U251 cells, and neuroblastoma SH-SY5Y cells. Co-culture of SH-SY5Y and U251 cells markedly enhanced claudin-5 and VE-cadherin expression in hCMEC/D3 cells, accompanied by increased transendothelial electrical resistance and decreased permeability. Conditioned medium (CM) from SH-SY5Y cells (S-CM), U251 cells (U-CM), and co-culture of SH-SY5Y and U251 cells (US-CM) also promoted claudin-5 and VE-cadherin expression. Glial cell line-derived neurotrophic factor (GDNF) levels in S-CM and US-CM were significantly higher than CMs from hCMEC/D3 and U-CM. Both GDNF and US-CM upregulated claudin-5 and VE-cadherin expression, which were attenuated by anti-GDNF antibody and GDNF signaling inhibitors. GDNF increased claudin-5 expression via the PI3K/AKT/FOXO1 and MAPK/ERK pathways. Meanwhile, GDNF promoted VE-cadherin expression by activating PI3K/AKT/ETS1 and MAPK/ERK/ETS1 signaling. The roles of GDNF in BBB integrity were validated using brain-specific *Gdnf* silencing mice. The developed triple co-culture BBB model was successfully applied to predict BBB permeability. In conclusion, neurons enhance BBB integrity by upregulating claudin-5 and VE-cadherin expression through GDNF secretion and established triple co-culture BBB model may be used to predict drugs’ BBB permeability.

**Impact Statement:** Neurons and astrocytes enhance the integrity of BBB by releasing GDNF. The released GDNF upregulated claudin-5 and VE-cadherin expression by the activation of PI3K/AKT and MAPK/ERK pathways.

## Introduction

As a dynamic interface between the blood circulatory system and the central nervous system (CNS), the blood-brain barrier (BBB) maintains homeostasis and normal function of CNS by strictly regulating material exchange between the blood and brain (Palmiotti et al., 2014). The maintenance of BBB is mainly attributed to the expression of tight junctions (TJ) and adherent junctions (AJ) between adjacent brain endothelial cells as well as a variety of drug transporters. However, as a double-edged sword that protects CNS function, BBB also restricts the transport of some drugs from blood to brain, leading to poor CNS therapeutic effects and even CNS treatment failure (Banks, 2016).

Several *in silico, in vitro, in situ,* and *in vivo* methods have been developed to assess the permeability of drugs across BBB, but each method has its limitations (Hanafy et al., 2021). The *in situ* brain perfusion (ISBP) is considered the “gold standard” for assessing BBB permeability but there exist limits related to animal ethics. Moreover, ISBP is not suitable for human or high-throughput workflows. Thus, a suitable, accurate, and high-throughput *in vitro* BBB model is required to predict the permeability of drug candidates through BBB.

BBB is formed by neurovascular units (NVU), composed of neural (neurons, microglia, and astrocytes) and vascular components (vascular endothelial cells, pericytes, and vascular smooth muscle cells) (Potjewyd et al., 2018). The constant crosstalk and interactions among these cells contribute to the structural and signaling-based regulation of transcellular and paracellular transport, control of BBB permeability, and regulation of cerebral circulation (Arvanitis et al., 2020; Muoio et al., 2014; Potjewyd et al., 2018). Although brain microvascular endothelial cells (BMECs) are commonly utilized as an *in vitro* BBB model to assess drug permeability across BBB, *in vitro* mono-culture of BMECs tends to lose some unique characteristics without the support of other cell types. To overcome the drawbacks of mono-culture BBB models, a range of multicellular co-culture BBB models co-cultured with other NVU elements (such as astrocytes or pericytes) have been developed. Astrocytes are the most abundant glial cell type in brain (Clasadonte et al., 2017). Their terminal feet cover >80% of the surface of capillaries to form interdigitating coverage without slits (Mathiisen et al., 2010). In addition, astrocytes also contain several proteins related to the tight binding of the basement membrane. The *in vitro* multicellular co-culture BBB models composed of astrocytes, pericytes, and endothelial cells are considered to mimic the vascular structure and offer paracellular tightness in BBB (Ito et al., 2019; Nakagawa et al., 2009; Watanabe et al., 2021).

Neurons also play a crucial role in the NVU and may be involved in the regulation of BBB function. The maturation of BBB in mice considerably overlaps with the establishment of neuronal activity (Biswas et al., 2020). A study revealed that the conditioned medium collected from primary rat neurons also attenuated cell death caused by glucose-oxygen-serum deprivation (Lin et al., 2016). Furthermore, the primary rat neurons were reported to affect the differentiation and formation of BMECs (Savettieri et al., 2000; Schiera G. et al., 2003; Xue et al., 2013). Recent research reported that, compared with the double co-culture of hCMEC/D3 and 1321N1 (human astrocytoma cells), the triple co-culture of hCMEC/D3, 1321N1, and human neuroblastoma SH-SY5Y cells exhibited a higher transendothelial electrical resistance (TEER) (Barberio et al., 2022). A similar report showed that co-cultivation of RBE4.B (rat brain capillary endothelial cells) and neurons resulted in the lower permeability of [^3^H] sucrose than RBE4.B cells grown alone (Schiera Gabriella. et al., 2005). All these studies indicate that neurons, as the elements of the NVU, may be tightly connected to the formation and maintenance of BBB functions.

The aims of the study were: 1) to establish and characterize an *in vitro* triple co-culture BBB model consisting of human brain endothelial cells (hCMEC/D3), human astrocytoma cells (U251), and human neuroblastoma cells (SH-SY5Y); 2) to investigate whether neurons were involved in the formation and maintenance of BBB integrity and explore their underlying mechanisms. The integrity of BBB was evaluated by quantifying the leakage of both fluorescein and FITC-Dextran 3–5 kDa (FITC-Dex); 3) to validate the *in vitro* results through *in vivo* experiments. Finally, the *in vivo*/*in vitro* correlation assay (IVIVC) was analyzed to prove that the triple co-culture BBB model could better predict the BBB penetration of CNS drugs compared with the mono-culture BBB model.

## Result

### Establishment and characterization of the *in vitro* triple co-culture BBB model

Four types of BBB *in vitro* models were established to compare the contributions of U251 and SH-SY5Y cells to hCMEC/D3 cells (Figure 1A). TEER values were measured during the co-culture (Figure 1B). TEER values of the four *in vitro* BBB models gradually increased until day 6. On day 7, the TEER values showed a decreasing trend. Thus, six-day co-culture period was used for subsequent experiments. The highest TEER values were observed in the triple co-culture BBB model, followed by double co-culture with U251 cells and double co-culture with SH-SY5Y cells BBB models, hCMEC/D3 cells mono-culture BBB model showed the lowest TEER values. We also measured the TEER values of U251 monolayer cells, and the results showed that U251 cells themselves also contributed to the physical barrier of the model (Figure 1C). The apparent permeability coefficient (*P_app_*) values of fluorescein and FITC-Dex were measured to characterize the integrity of four BBB models (Figure 1D, 1E). Consistent with the TEER values, the triple co-culture BBB model showed the lowest permeability of fluorescein and FITC-Dex, followed by double co-culture with U251 cells and double co-culture with SH-SY5Y cells BBB models. It was noticed that monolayer of U251 cells itself also worked as a barrier, preventing the leakage of permeability markers, which may explain why the permeability of FITC-Dex in double co-culture model with U251 cells is lower than that in double co-culture model with SH-SY5Y cells (Figure 1F). Co-culture with SH-SY5Y, U251, and U251 + SH-SY5Y cells also enhanced the proliferation of hCMEC/D3 cells. Moreover, the promoting effect of SH-SY5Y cells was stronger than that of U251 cells (Figure 1G-1I). Furthermore, hCMEC/D3 cells were incubated with basic fibroblast growth factor (bFGF), which promotes cell proliferation without affecting both claudin-5 and VE-cadherin expression (Figure 2F). The results showed that incubation with bFGF increased cell proliferation and reduced permeabilities of fluorescein and FITC-Dex across hCMEC/D3 cell monolayer. However, the permeability reduction was less than that by double co-culture with U251 cells or triple co-culture. These results inferred that contribution of cell proliferation to the barrier function of hCMEC/D3 was minor (Figure 1—figure supplement 1).

**Figure 1.**
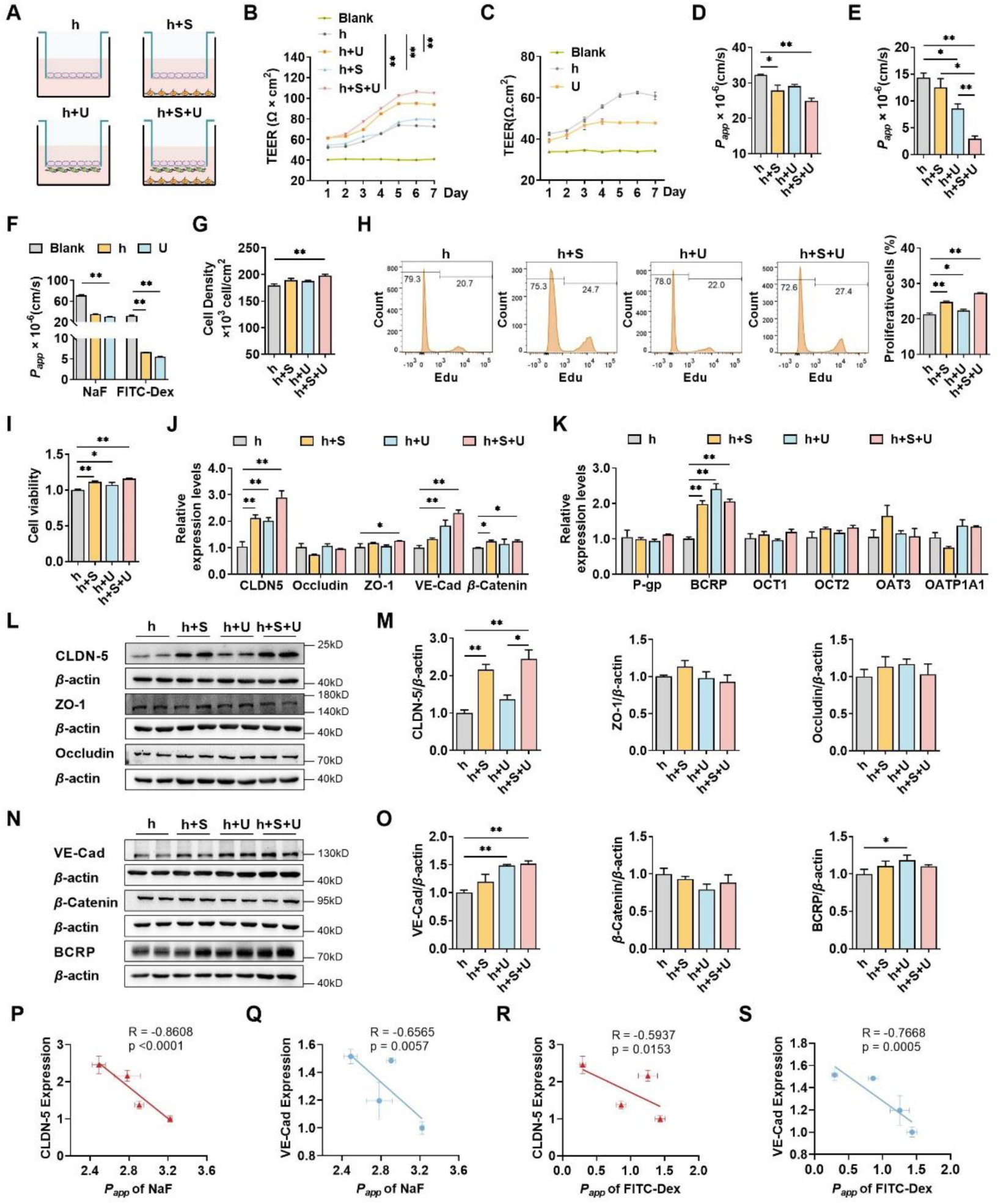
The effects of co-culture with U251 and/or SH-SY5Y cells on the integrity of hCMEC/D3 and BBB function. (**A**): Four different types of BBB models were prepared from hCMEC/D3 cells (h), SH-SY5Y cells (S), and U251 cells (U). (**B**): The transendothelial electrical resistance (TEER) of four models, and the TEER values in day 6 were compared. Blank: no cells. Four biological replicates per group. (**C**): The TEER of hCMEC/D3 and U251 cells monolayer. Four biological replicates per group. (**D** and **E**): The apparent permeability coefficient (*P_app_*, × 10^-6^ cm/s) of fluorescein (NaF) and FITC-Dextran 3–5 kDa (FITC-Dex) of four BBB models. Four biological replicates per group. (**F**): The *P_app_* (× 10^-6^ cm/s) of NaF and FITC-Dex across the blank inserts, and hCMEC/D3 or U251 mono-culture models. Four biological replicates per group. (**G** and **H**): The cell density (G), EdU incorporation (H) of hCMEC/D3 cells after mono/co-culturing. Three biological replicates per group. (**I**): Cell viability of hCMEC/D3 cells after mono/co-culturing. Four biological replicates per group. (**J** and **K**): The mRNA levels of tight junction proteins, adherent junction proteins, and transporters. Four biological replicates per group. (**L-O**): The protein expression levels of claudin-5 (CLDN-5), ZO-1, occluding (L and M), VE-cadherin (VE-Cad), *β*-catenin, and BCRP (N and O) in hCMEC/D3 cells. Four biological replicates per group. (**P** and **Q**): The correlations between the *P_app_* (× 10^-5^ cm/s) of NaF and claudin-5 expression (P), or VE-cadherin expression (Q). (**R** and **S**): The correlation between *P_app_* (× 10^-5^ cm/s) of FITC-Dex and claudin-5 expression (R), or VE-cadherin expression (S). The above data are shown as the mean ± SEM. For J and K, two technical replicates per biological replicate. One technical replicate per biological replicate for the rest. * *p* < 0.05; ** *p* < 0.01 by one-way ANOVA test followed by Fisher’s LSD test, Welch’s ANOVA test, or Kruskal-Wallis test. The simple linear regression analysis was used to examine the presence of a linear relationship between two variables. **Figure 1-Source data1** The western blot raw images in Figure 1 **Figure 1-Source data2** The labeled western blot images in Figure 1 **Figure 1-Source data3** Excel file containing summary data and data analysis of Figure 1

**Figure 2.**
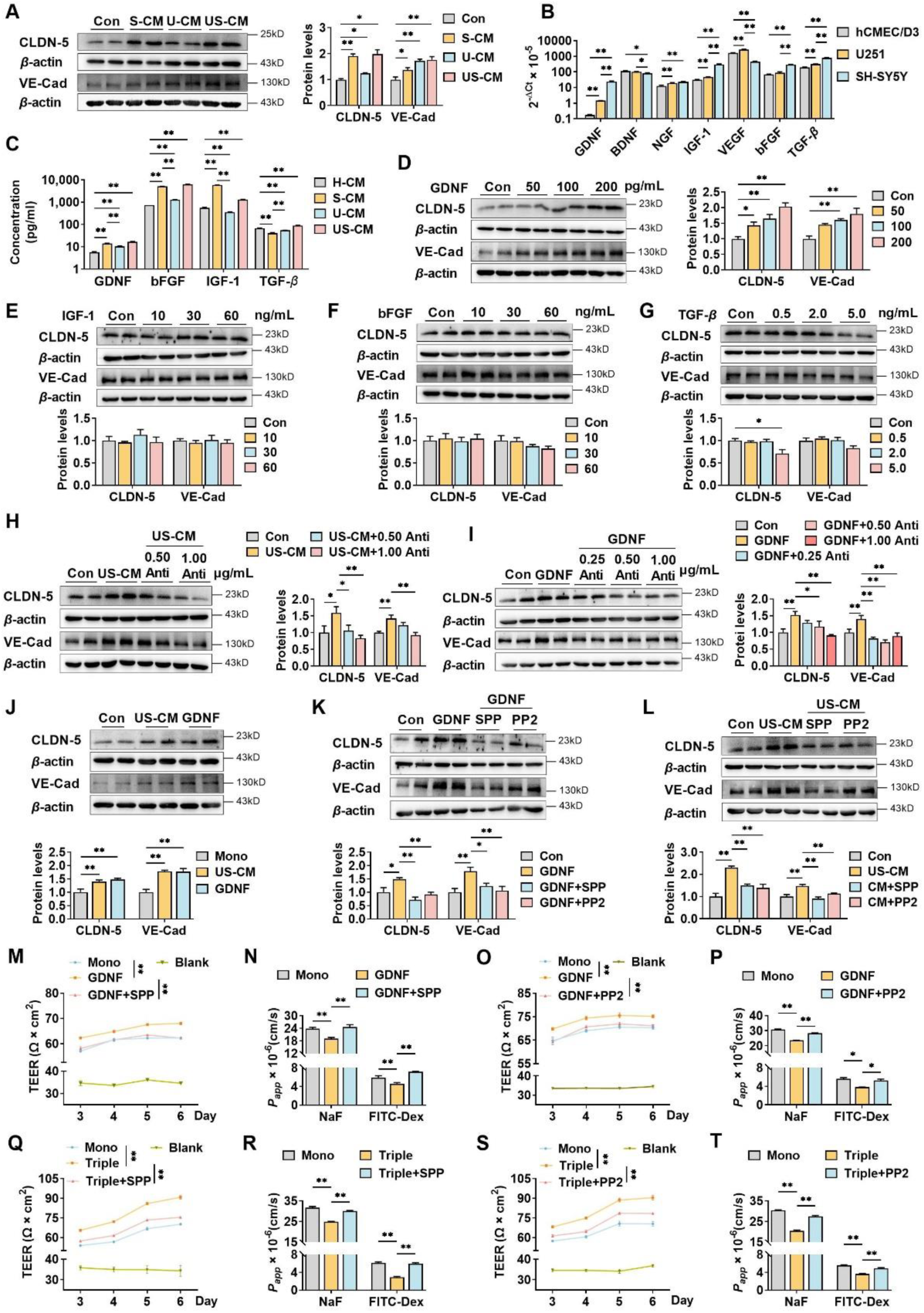
Neurons and astrocytes upregulated claudin-5 and VE-cadherin expression in hCMEC/D3 cells due to GDNF secretion. (**A**): Effects of conditioned medium (CM) on claudin-5 and VE-cadherin expression. Con: the normal medium; S-CM: the CM from SH-SY5Y cells; U-CM: the CM from U251 cells; US-CM: the CM from SH-SY5Y cells co-culture with U251 cells. (**B**): The mRNA expression levels of neurotrophic factors in hCMEC/D3, U251, and SH-SY5Y cells. (**C**): Concentrations of GDNF, bFGF, IGF-1, and TGF-*β* in the CMs. H-CM: the CM from hCMEC/D3 cells. (**D**-**G**): Effects of GDNF (D), IGF-1 (E), bFGF (F), and TGF-*β* (G) on the expression of claudin-5 and VE-cadherin. The dosages have been marked in the figure. (**H** and **I**): Effects of anti-GDNF antibody on the upregulation of claudin-5 and VE-cadherin expression induced by US-CM (H) or 200 pg/mL GDNF (I). (**J**): Effects of 200 pg/mL GDNF and US-CM on claudin-5 and VE-cadherin expression in primary rat brain microvascular endothelial cells. (**K** and **L**): Effects of 3 μM RET tyrosine kinase inhibitor SSP-86 (SPP), and 5 μM Src family kinases inhibitor PP2 on the upregulation of claudin-5 and VE-cadherin induced by 200 pg/mL GDNF (K) and US-CM (L). (**M** and **N**): Effects of SPP on the TEER on day 6 (M), the permeability of NaF, and FITC-Dex (N) of the hCMEC/D3 mono-culture BBB model treating 200 pg/mL GDNF. (**O** and **P**): Effects of PP2 on the TEER on day 6 (O), the permeability of NaF, and FITC-Dex (P) of the hCMEC/D3 mono-culture BBB model treating 200 pg/mL GDNF. (**Q** and **R**): Effects of SPP on the TEER on day 6 (Q), the permeability of NaF, and FITC-Dex (R) of the triple co-culture BBB model. (**S** and **T**): Effects of PP2 on the TEER on day 6 (S), the permeability of NaF, and FITC-Dex (T) of the triple co-culture BBB model. The above data are shown as the mean ± SEM. Four biological replicates per group. For B and C, two technical replicates per biological replicate. One technical replicate per biological replicate for the rest. * *p* < 0.05; ** *p* < 0.01 by one-way ANOVA test followed by Fisher’s LSD test, Welch’s ANOVA test, or Kruskal-Wallis test. **Figure 2-Source data1** The western blot raw images in Figure 2 **Figure 2-Source data2** The labeled western blot images in Figure 2 **Figure 2-Source data3** Excel file containing summary data and data analysis of Figure 2

The paracellular barrier of BBB is also associated with TJs, AJs, and transporters (Abbott, 2013). The mRNA levels of TJs (claudin-5, ZO-1, and occludin), AJs (VE-cadherin and *β*-catenin), and transporters (P-gp, BCRP, OCT-1, OCT-2, OAT-3, and OATP1A1) in hCMEC/D3 cells from the four BBB models were analyzed using qPCR (Figure 1J and 1K). Compared with hCMEC/D3 cell mono-culture model, double co-culture with SH-SY5Y, double co-culture with U251, and triple co-culture BBB models showed markedly increases in claudin-5, VE-cadherin, *β*-catenin, and BCRP mRNA expression. Expression of corresponding proteins was measured using Western blot (Figure 1L-1O). Notably increased claudin-5 expression was detected in double co-culture with SH-SY5Y cells and triple co-culture BBB models, while VE-cadherin expression was markedly increased in double co-culture with U251 cells and triple co-culture BBB models. Expression levels of other TJ proteins (ZO-1 and occludin) and AJ protein (*β*-catenin) were unaltered. The expression of BCRP was slightly affected by co-cultivation with U251 cells. Significant negative correlations were found between *P_app_* values of fluorescein and the expression of claudin-5 or VE-cadherin. *P_app_* values of FITC-Dex were also negatively correlated to the expression levels of claudin-5 or VE-cadherin (Figure 1P-1S). These results indicate that the decreased permeability of fluorescein and FITC-Dex mainly results from the upregulated expression of both claudin-5 and VE-cadherin.

### Neurons and astrocytes upregulated the expression of claudin-5 and VE-cadherin by GDNF secretion

The hCMEC/D3 cells did not direct contact with U251 or SH-SY5Y cells in the double co-culture and triple co-culture BBB models, indicating that cell-cell interaction between U251, SH-SY5Y, and hCMEC/D3 cells relied on secreted active factors. To test this hypothesis, the effects of conditioned medium (CM) from SH-SY5Y cells (S-CM), U251 cells (U-CM), and co-culture of SH-SY5Y and U251 cells (US-CM) on the expression of claudin-5 and VE-cadherin in hCMEC/D3 cells were analyzed (Figure 2A). Both S-CM, U-CM, and US-CM markedly increased the expression of claudin-5 and VE-cadherin. US-CM showed the strongest induction effects on claudin-5 and VE-cadherin.

To investigate which cytokines were involved in the promotion of hCMEC/D3 cell integrity by U251 and SH-SY5Y cells, the mRNA expression levels of various cytokines in these three types of cells were compared. The results showed that U251 or SH-SY5Y cells exhibited significantly higher expression levels of GDNF, nerve growth factor (NGF), insulin-like growth factor-1 (IGF-1), vascular endothelial growth factor (VEGF), bFGF, and transforming growth factor-*β* (TGF-*β*) compared to hCMEC/D3 cells (Figure 2B). Furthermore, the mRNA expression of GDNF, IGF-1, TGF-*β*, and bFGF in SH-SY5Y cells was higher than those in U251 cells In these cytokines, GDNF (Dong et al., 2018; Igarashi et al., 1999; Shimizu et al., 2012), bFGF (Shimizu et al., 2011; Wang et al., 2016), IGF-1 (Ji-Ae K et al., 2009; Nowrangi et al., 2019), and TGF-*β* (Fu et al., 2021) have been reported to promote BBB integrity. Thus, the concentrations of GDNF, bFGF, IGF-1, and TGF-*β* in the CMs were measured (Figure 2C). The results showed that levels of GDNF, bFGF, and IGF-1 in S-CM and US-CM were significantly higher than CMs from hCMEC/D3 cells (H-CM) and U-CM, but levels of TGF-*β* in S-CM and U-CM were lower than those in H-CM. Interestingly, the level of IGF-1 in US-CM was remarkably lower than that in S-CM, indicating that U251 cells suppressed IGF-1 secretion from SH-SY5Y. The effects of GDNF, bFGF, IGF-1, and TGF-*β* on the expression of claudin-5 and VE-cadherin were investigated (Figure 2D-2G). Among the four tested neurotrophic factors, only GDNF induced the expression of claudin-5 and VE-cadherin in a concentration-dependent manner (Figure 2D). In contrast, a high level (5 ng/mL) of TGF-*β* slightly downregulated claudin-5 expression (Figure 2G). These results demonstrate that upregulation of claudin-5 and VE-cadherin expression by US-CM are attributed to secreted GDNF.

To provide additional verification of the deduction, anti-GDNF antibody was used to neutralize exogenous and endogenous GDNF in culture medium. Consistent with our expectation, the anti-GDNF antibody concentration-dependent reversed the US-CM-induced claudin-5 and VE-cadherin expression (Figure 2H). Furthermore, GDNF-induced upregulation of claudin-5 and VE-cadherin expression was also reversed by the anti-GDNF antibody (Figure 2I). The induction effects of US-CM and GDNF on the claudin-5 and VE-cadherin expression in hCMEC/D3 cells were also confirmed in primary rat brain microvascular endothelial cells (Figure 2J)GDNF forms a heterohexameric complex with two GFR*α*1 molecules and two RET receptors to activate the GDNF-GFR*α*1-RET signaling (Fielder et al., 2018). The RET receptor tyrosine kinase inhibitor SPP-86 (Bhallamudi et al., 2021) and Src-type kinase inhibitor PP2 (Morita et al., 2006) were used to further investigate whether GDNF upregulated the expression of claudin-5 and VE-cadherin in hCMEC/D3 cells by activating the GDNF-GFR*α*1-RET signaling pathway. Both SPP-86 and PP2 markedly attenuated claudin-5 and VE-cadherin expression induced by GDNF (Figure 2K) and US-CM (Figure 2L).

The contributions of GDNF-induced claudin-5 and VE-cadherin expression to TEER and permeability were investigated using hCMEC/D3 cells mono-culture BBB model. GDNF significantly increased TEER values (Figure 2M, 2O) and decreased the permeability of fluorescein and FITC-Dex (Figure 2N, 2P), which were almost abolished by SPP-86 or PP2. Furthermore, treatment with SPP-86 or PP2 completely reversed the increased TEER values (Figure 2Q, 2S) and decreased permeability of fluorescein and FITC-Dex (Figure. 2R, 2T) in the triple co-culture BBB model. These results indicate that neurons but also astrocytes upregulate claudin-5 and VE-cadherin expression in hCMEC/D3 cells by secreting GDNF. Subsequently, GDNF induces claudin-5 and VE-cadherin expression by activating GDNF-GFR*α*1-RET signaling.

### GDNF induced claudin-5 and VE-cadherin expression of hCMEC/D3 by activating the PI3K/AKT and MAPK/ERK pathways

GDNF exerts its biological activities by activating several signaling pathways, including the phosphatidylinositol-3-kinase (PI3K)/ protein kinase B (AKT), mitogen-activated protein kinase (MAPK)/ extracellular regulated kinase (ERK), MAPK/ c-Jun N-terminal kinase (JNK), and MAPK/ p38 pathways (Fielder et al., 2018). The effects of the PI3K/AKT, MAPK/ERK, MAPK/JNK, and MAPK/p38 pathway inhibitors LY294002 (Figure 3A), U0126 (Figure 3B), SP600125 (Figure 3C), and SB203580 (Figure 3D), respectively, on GDNF-induced claudin-5 and VE-cadherin expression in hCMEC/D3 cells were investigated. GDNF increased claudin-5 and VE-cadherin expression, accompanied by the phosphorylation of AKT (p-AKT) and ERK (p-ERK). However, it did not stimulate the phosphorylation of JNK and p38. SPP-86, LY29002, and U0126 significantly suppressed GDNF-induced claudin-5 and VE-cadherin expression, while SP600125 and SB203580 had almost no effect on GDNF-induced claudin-5 and VE-cadherin expression. GDNF-induced phosphorylation of AKT and ERK was also markedly attenuated by the anti-GDNF antibody (Figure 3E). Similarly, US-CM remarkably upregulated the expression of claudin-5, VE-cadherin, p-AKT, and p-ERK, which were also markedly reversed by SPP-86, LY29002, U0126, or anti-GDNF antibody (Figure 3F-3J). These findings indicate that GDNF induces the expression of claudin-5 and VE-cadherin in hCMEC/D3 cells by activating both the PI3K/AKT and MAPK/ERK pathways.

**Figure 3.**
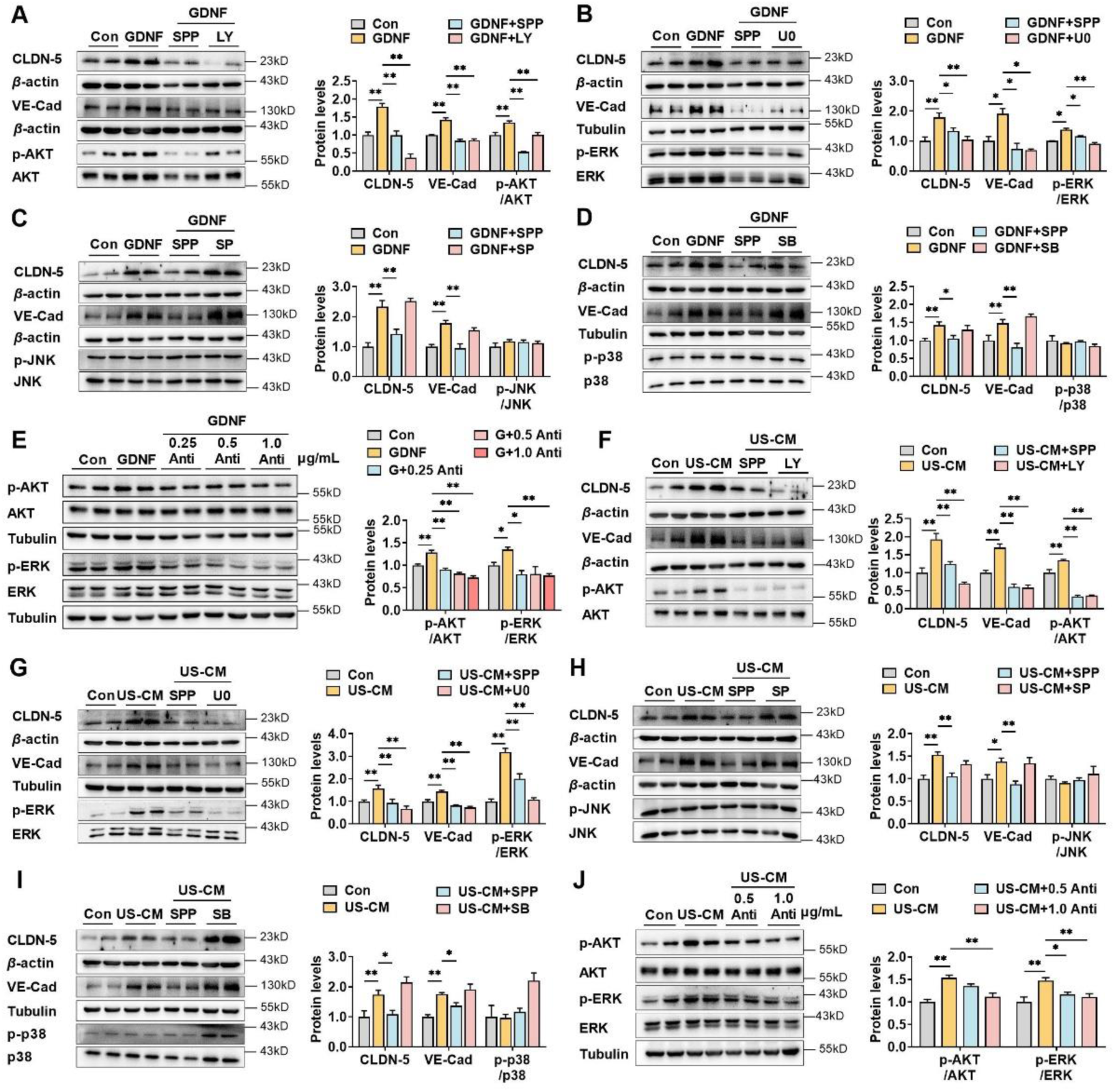
GDNF induced claudin-5 and VE-cadherin expression in hCMEC/D3 cells by activating the PI3K/AKT and MAPK/ERK signaling. (**A**): Effects of 3 μM LY294002 (LY) on the levels of claudin-5, VE-cadherin, and p-AKT/AKT in hCMEC/D3 cells stimulated by 200 pg/mL GDNF. (**B**): Effects of 2 μM U0126 (U0) on the levels of claudin-5, VE-cadherin, and p-ERK/ERK in hCMEC/D3 cells stimulated by 200 pg/mL GDNF. (**C**): Effects of 5 μM SP600125 (SP) on the levels of claudin-5, VE-cadherin, and p-JNK/JNK in hCMEC/D3 cells stimulated by 200 pg/mL GDNF. (**D**): Effects of 2 μM SB203580 (SB) on the levels of claudin-5, VE-cadherin, and p-p38/p38 in hCMEC/D3 cells stimulated by 200 pg/mL GDNF. (**E**): Effects of anti-GDNF antibody on the GDNF-induced p-AKT/AKT and p-ERK/ERK ratios. (**F**): Effects of 3 μM LY on the levels of claudin-5, VE-cadherin, and p-AKT/AKT in hCMEC/D3 cells stimulated by US-CM. (**G**): Effects of 2 μM U0 on the levels of claudin-5, VE-cadherin, and p-ERK/ERK in hCMEC/D3 cells stimulated by US-CM. (**H**): Effects of 5 μM SP on the levels of claudin-5, VE-cadherin, and p-JNK/JNK in hCMEC/D3 cells stimulated by US-CM. (**I**): Effects of 2 μM SB on the levels of claudin-5, VE-cadherin, and p-p38/p38 in hCMEC/D3 cells stimulated by 2 US-CM. (**J**): Effects of anti-GDNF antibody on the US-CM-induced p-AKT/AKT and p-ERK/ERK ratios. The above data are shown as the mean ± SEM. Four biological replicates per group. One technical replicate for each biological replicate. * *p* < 0.05; ** *p* < 0.01 by one-way ANOVA test followed by Fisher’s LSD test or Welch’s ANOVA test. **Figure 3-Source data1** The western blot raw images in Figure 3 **Figure 3-Source data2** The labeled western blot images in Figure 3 **Figure 3-Source data3** Excel file containing summary data and data analysis of Figure 3

### GDNF upregulated the claudin-5 expression in hCMEC/D3 cells by activating the PI3K/AKT/FOXO1 pathway

Claudin-5 is negatively regulated by the transcriptional repressor forkhead box O1 (FOXO1) (Beard et al., 2020). FOXO1 is also an important target of PI3K/AKT signaling. FOXO1 phosphorylation results in FOXO1 accumulation in the cytoplasm (Zhang et al., 2011) and lowers its level in the nucleus. Here, we investigated whether GDNF-induced claudin-5 expression is involved in FOXO1 nuclear exclusion. As shown in Figure 4A, both GDNF and US-CM significantly enhanced FOXO1 phosphorylation (p-FOXO1). Similarly, GDNF and US-CM increased the levels of phosphorylated and unphosphorylated FOXO1 in the cytoplasm and decreased the levels of nuclear FOXO1 (Figure 4B). Whether FOXO1 was involved in the GDNF-induced regulation of claudin-5 and VE-cadherin expression was investigated in hCMEC/D3 cells transfected with *FOXO1* siRNA. Silencing *FOXO1* significantly decreased FOXO1 levels in both whole-cell lysates and nucleus of hCMEC/D3 cells (Figure 4C), demonstrating *FOXO1* silencing efficacy. Consistent with our expectation, silencing *FOXO1* upregulated the expression of claudin-5 rather than VE-cadherin expression (Figure 4D). In contrast, high levels of FOXO1 were observed in both whole-cell lysates and nucleus of hCMEC/D3 cells that were transfected with plasmids containing *FOXO1*. Meanwhile, FOXO1 overexpression resulted in a decrease in both basal and GDNF-induced claudin-5 expression (Figure 4E), consistent with the known role of FOXO1 on claudin-5 expression.

**Figure 4.**
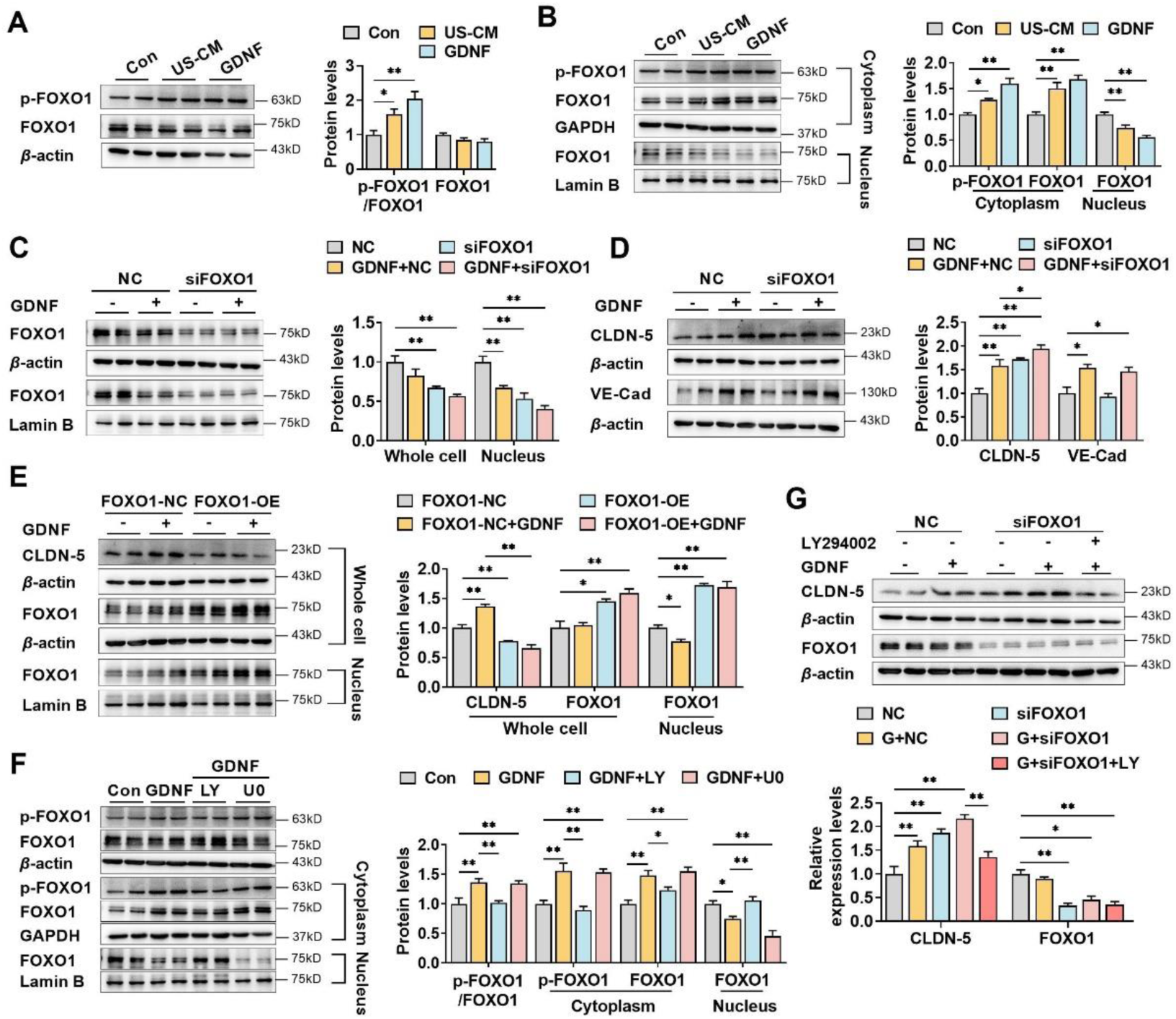
GDNF induced the claudin-5 expression in hCMEC/D3 cells by activating the PI3K/AKT/FOXO1 pathway. (**A** and **B**): Effects of US-CM and GDNF on the phosphorylated FOXO1 (p-FOXO1)/FOXO1 ratio, total FOXO1 expression (A), cytoplasmic p-FOXO1, cytoplasmic FOXO1, and nuclear FOXO1 expression (B). (**C** and **D**): The expression levels of total and nuclear FOXO1 (C), claudin-5, and VE-cadherin (D) in hCMEC/D3 cells transfected with *FOXO1* siRNA (siFOXO1). NC: negative control. (**E**): Effects of *FOXO1* overexpression (FOXO1-OE) and GDNF on the expression levels of claudin-5, total FOXO1, and nuclear FOXO1. FOXO1-NC: negative control plasmids. (**F**): Effects of LY and U0 on GDNF-induced alterations of total p-FOXO1/FOXO1 ratio, cytoplasmic p-FOXO1, cytoplasmic FOXO1, and nuclear FOXO1 expression. (**G**): Effects of LY on the claudin-5 expression upregulated by siFOXO1. The above data are shown as the mean ± SEM. Four biological replicates per group. One technical replicate for each biological replicate. * *p* < 0.05; ** *p* < 0.01 by one-way ANOVA test followed by Fisher’s LSD test, Welch’s ANOVA test, or Kruskal-Wallis test. **Figure 4-Source data1** The western blot raw images in Figure 4 **Figure 4-Source data2** The labeled western blot images in Figure 4 **Figure 4-Source data3** Excel file containing summary data and data analysis of Figure 4

Several reports have demonstrated that nuclear localization of FOXO1 is modulated by multiple pathways, including the PI3K/AKT (Tang et al., 1999) and MAPK/ERK (Asada et al., 2007) pathways. The effects of LY294002 and U0126 on GDNF-induced FOXO1 phosphorylation were measured in hCMEC/D3 cells.

LY294002, but not U0126, significantly reversed the GDNF-induced alterations in total p-FOXO1, cytoplasmic p-FOXO1, cytoplasmic FOXO1, and nuclear FOXO1 (Figure 4F). Furthermore, LY294002 reversed the upregulation of claudin-5 expression induced by *FOXO1* siRNA and GDNF (Figure 4G).

It was reported that VE-cadherin also upregulates claudin-5 via inhibiting FOXO1 activities (Taddei et al., 2008). Effect of VE-cadherin on claudin-5 was studied in hCMEC/D3 cells silencing VE-cadherin. It was not consistent with Taddei et al. that silencing VE-cadherin only slightly decreased the mRNA level of claudin-5 without significant difference. Furthermore, basal and GDNF-induced claudin-5 protein levels were unaltered by silencing VE-cadherin (Figure 4—figure supplement 1). Thus, the roles of VE-cadherin in regulation of claudin-5 in BBB should be further investigated.

### GDNF upregulated VE-cadherin expression in hCMEC/D3 cells by activating the PI3K/AKT/ETS1 and MAPK/ERK/ETS1 signaling pathways

E26 oncogene homolog 1 (ETS1) is a transcription factor that binds to the ETS-binding site located in the proximal region of the VE-cadherin promoter, hence regulating the expression of VE-cadherin (Lelièvre et al., 2000; Luo et al., 2022). Activation of the PI3K/AKT (He et al., 2023; Hui et al., 2018) and MAPK/ERK (Watanabe et al., 2004) pathways was reported to upregulate the expression of ETS1. The previous results showed that GDNF induced VE-cadherin and claudin-5 expression in hCMEC/D3 cells by activating the PI3K/AKT and MAPK/ERK pathways. Therefore, we hypothesized that GDNF modulated ETS1 levels to promote VE-Cadherin and claudin-5 expression through PI3K/AKT and MAPK/ERK signaling pathways. Both US-CM and GDNF significantly increased total (Figure 5A) and nuclear ETS1 expression (Figure 5B). LY294002 and U0126 markedly attenuated GDNF-induced total (Figure 5C) and nuclear ETS1 expression (Figure 5D). To further confirm the involvement of the PI3K/AKT/ETS1 and MAPK/ERK/ETS1 pathways in GDNF-induced VE-cadherin expression, ETS1 in hCMEC/D3 cells was knocked down using *ETS1* siRNA*. ETS1* silencing saliently declined the expression levels of total (Figure 5E) and nuclear (Figure 5F) ETS1 in hCMEC/D3 cells, demonstrating silencing efficacy. In *ETS1* silencing hCMEC/D3 cells, GDNF no longer induced the expression of total (Figure 5E) and nuclear (Figure 5F) ETS1. Moreover, *ETS1* silencing substantially downregulated VE-cadherin expression and attenuated GDNF-induced VE-cadherin expression (Figure 5G and 5H), while having minimal impact on both basal and GDNF-induced claudin-5 expression (Figure 5G and 5I).

**Figure 5.**
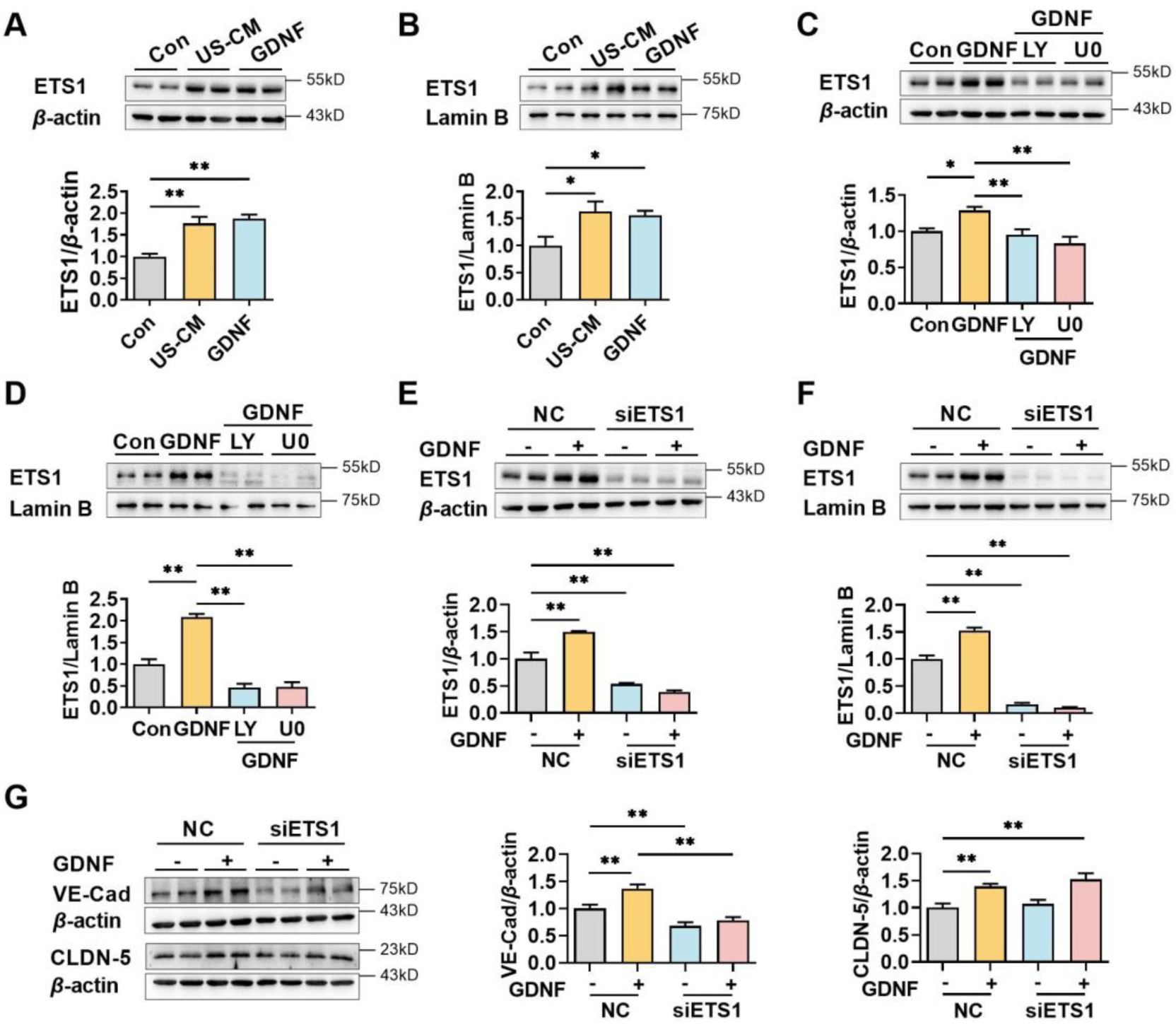
GDNF induced VE-cadherin expression in hCMEC/D3 cells by activating the PI3K/AKT/ETS1 and MAPK/ERK/ETS1 pathways. (**A** and **B**): Effects of US-CM and GDNF on total (A) and nuclear (B) ETS1 expression. (**C** and **D**): Effects of LY and U0 on 200 pg/mL GDNF-induced total (C) and nuclear (D) ETS1 expression. (**E** and **F**): Expression levels of total (E) and the nuclear ETS1 (F) in hCMEC/D3 cells after knocking down *ETS1* with siRNA (siETS1). (**G**-**I**): Effects of GDNF and siETS1 on the expression of VE-cadherin and claudin-5. The above data are shown as the mean ± SEM. Four biological replicates per group. One technical replicate for each biological replicate. * *p* < 0.05; ** *p* < 0.01 by one-way ANOVA test followed by Fisher’s LSD test. **Figure 5-Source data1** The western blot raw images in Figure 5 **Figure 5-Source data2** The labeled western blot images in Figure 5 **Figure 5-Source data3** Excel file containing summary data and data analysis of Figure 5

### Brain GDNF deficiency increased BBB permeability partly due to the impairment of claudin-5 and VE-cadherin expression

To further demonstrate the positive effects of GDNF on BBB maintenance, GDNF in mice brains was knocked down via intracerebroventricular injection of AAV-PHP.eB packaged with *Gdnf* short hairpin RNA (shRNA) (Figure 6A). Knockdown efficiency was confirmed through Western blotting (Figure 6B). Consistent with *in vitro* results, GDNF knockdown greatly downregulated claudin-5 and VE-cadherin expression in the mice brains (Figure 6B). The integrity of BBB was assessed by examining the brain distributions of fluorescein and FITC-Dex. The results showed that specifically knocking down brain GDNF little affected plasma levels of fluorescein (Figure 6C) and FITC-Dex (Figure 6F), but significantly elevated the concentrations of fluorescein (Figure 6D) and FITC-Dex (Figure 6G) in the brains, leading to notable increases in the brain-to-plasma concentration ratios of two probes (Figure 6E and 6H). These alterations were consistent with the decline in claudin-5 and VE-cadherin expression. In addition, significant reductions in the levels of p-AKT (Figure 6I), p-ERK (Figure 6J), p-FOXO1 (Figure 6K), and ETS1 expression (Figure 6L) were observed in the brains of *Gdnf* knockdown mice.

**Figure 6.**
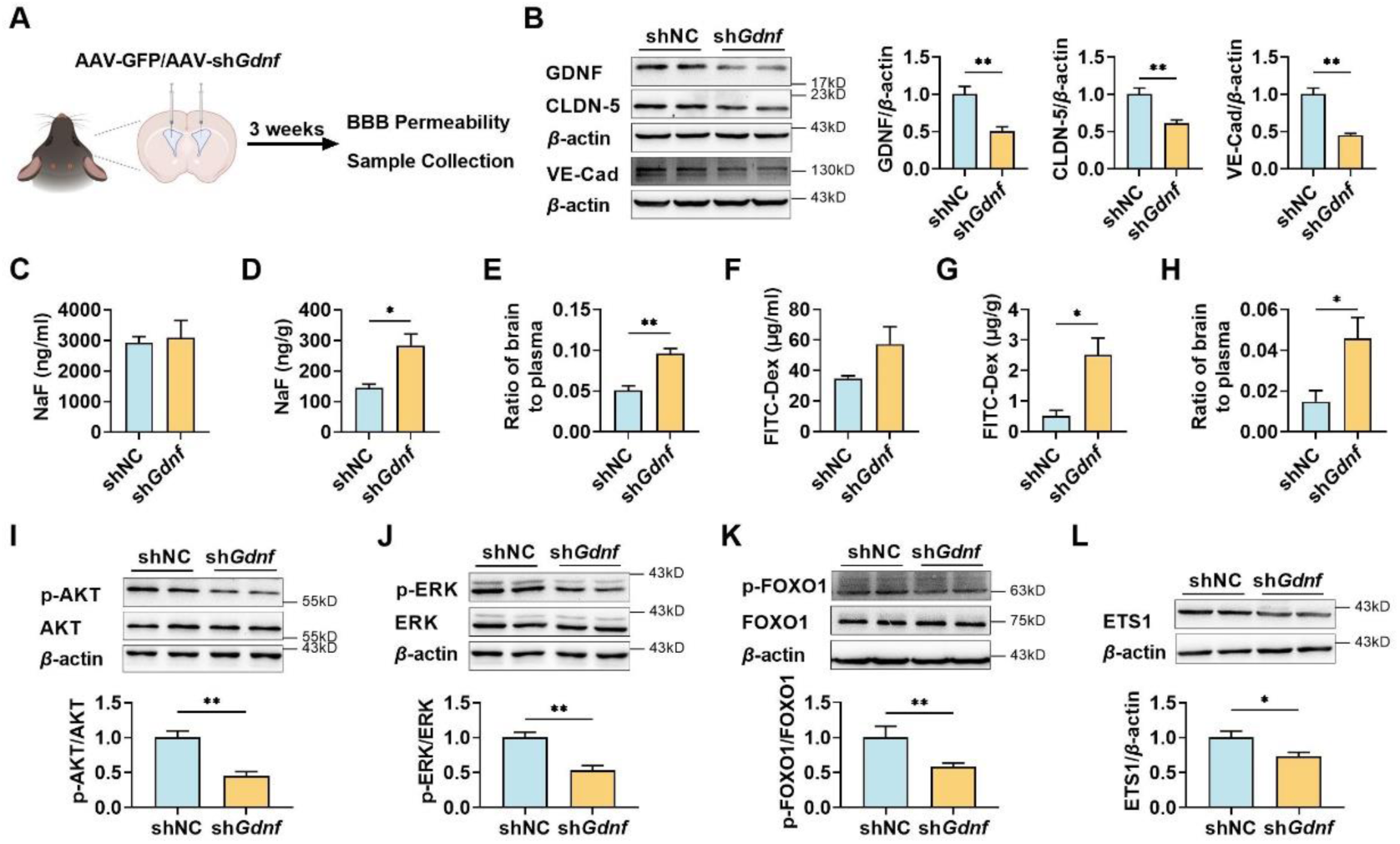
The deficiency of brain GDNF in mice increased the permeability of BBB and reduced claudin-5 and VE-cadherin expression in mice brains. (**A**): Experimental configuration of AAV-GFP (shNC) or AAV-sh*Gdnf* (sh*Gdnf*) intracerebroventricular injection. (**B**): Effects of brain-specific *Gdnf* silencing on the expression levels of GDNF, claudin-5, and VE-cadherin in the brains. (**C**-**E**): Effects of brain-specific *Gdnf* silencing on NaF levels in plasma (C), brain (D), and the ratio of brain to plasma (E). (**F**-**H**): Effects of brain-specific *Gdnf* silencing on FITC-Dex levels in plasma (F), brain (G), and the ratio of brain to plasma (H). (**I**-**K**): The expression ratios of p-AKT/AKT (I), p-ERK/ERK (J), and p-FOXO1/FOXO1 (K) in the brains of *Gdnf* silencing mice. (**L**): The expression level of ETS1 in the brains of *Gdnf* silencing mice. The above data are shown as the mean ± SEM. Six biological replicates per group. One technical replicate for each biological replicate. * *p* < 0.05; ** *p* < 0.01 by unpaired t-test, unpaired t-test with Welch’s correction, or Mann-Whitney test. **Figure 6-Source data1** The western blot raw images in Figure 6 **Figure 6-Source data2** The labeled western blot images in Figure 6 **Figure 6-Source data3** Excel file containing summary data and data analysis of Figure 6

### The triple co-culture BBB model better predicted the permeabilities of drugs across BBB

In this study, 18 drugs were utilized to further investigate the superiority of the triple co-culture BBB model over the hCMEC/D3 mono-culture BBB model. The *P_app_* of 18 drugs from the apical to the basolateral side based on the hCMEC/D3 mono-culture (*P_app, Mono_*) and triple co-culture (*P_app, Triple_*) BBB models are measured and listed in Table 1. The results showed that *P_app, Triple_*values of all tested drugs were lower than the *P_app, Mono_* values. Significant differences were observed in 14 out of 18 drugs. The predicted permeability coefficient-surface area product values (*PS*) of the tested drugs were respectively calculated based on their *P_app, Mono_*values (*PS _Pre, Mono_*), and *P_app, Triple_* values (*PS _Pre, Triple_*). The predicted *PS* values were further compared to their corresponding observations (*PS _obs_*). The results showed that the predictive accuracy of *PS _Pre, Triple_* was superior to *PS _Pre, Mono_*. Except for verapamil, amitriptyline, fluoxetine, and clozapine, the predicted *PS _Pre, Triple_* values of the other 14 drugs in the triple co-culture BBB models were within the 0.5–2 folds of *PS _obs_*(Figure 7B). However, in the hCMEC/D3 mono-culture BBB model, only 7 predicted *PS _Pre, Mono_* values were within the 0.5–2 folds range of their observations (Figure 7A).

**Figure 7.**
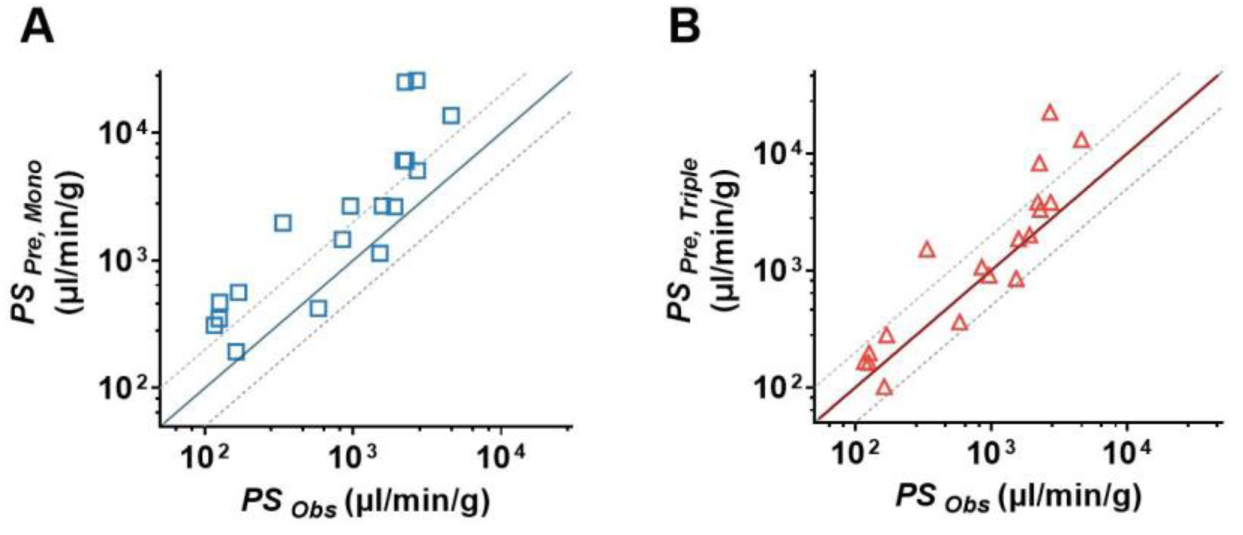
*In vitro*/*in vivo* correlation assay of BBB permeability. (**A**): The comparison of the estimated permeability coefficient-surface area product (*PS _Mono_*) recalculated from *P_app, Mono_* with the observed *in vivo PS* values (*PS _Obs_*). (**B**): The comparison of the estimated permeability coefficient-surface area product (*PS _Triple_*) recalculated from *P_app, Triple_* with the observed *in vivo PS* values (*PS _Obs_*). The solid line represents a perfect prediction, and the dashed lines represent the 0.5-2 folds of their observations. The *PS _Obs_* values were determined by *in situ* brain perfusion in rodents, which were collected from the literature.

**Table 1.**
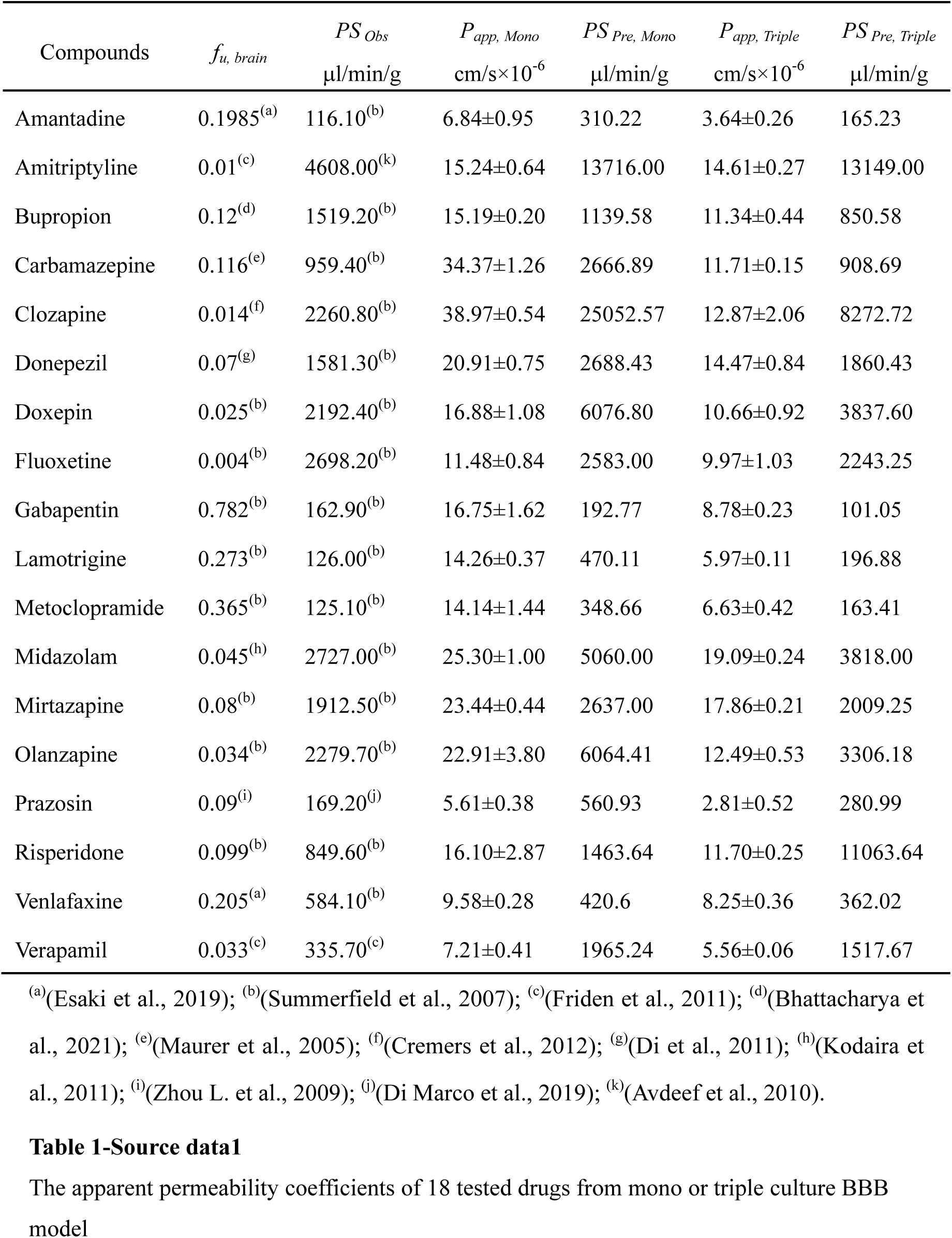
The unbound fraction in brain (*f_u, brain_*), the observed *PS _Obs_*, and the predicted *PS* (*PS _Pre_*), *P_app_* across the hCMEC/D3 mono-culture model (*P_app, Mono_*) and triple co-culture model (*P_app, Triple_*) of the tested drugs. One technical replicate of 4 biological replicates per group.

## Discussion

The main findings of the study were to successfully develop an *in vitro* triple co-culture BBB model consisting of hCMEC/D3, U251, and SH-SY5Y cells and to confirm the involvement of neurons in BBB maintenance as well as the possible mechanisms. Co-culture with U251 and/or SH-SY5Y cells markedly promoted the TEER of hCMEC/D3 cells and reduced the leakage of fluorescein and FITC-Dex due to the upregulation of claudin-5 and VE-cadherin expression.

The roles of claudin-5 and VE-cadherin in the maintenance of BBB function have been demonstrated (Dejana et al., 2008; Hashimoto et al., 2023; Li et al., 2018; Ohtsuki et al., 2007). It was reported that *Cld-5*–deficient mice exhibited BBB impairment, allowing the transport of small molecules (<800 D) across BBB (Nitta et al., 2003). In contrast, claudin-5 overexpression significantly restricted the permeability of inulin across the conditionally immortalized rat brain capillary endothelial cell monolayer (Ohtsuki et al., 2007). Moreover, the claudin-5 expression in hCMEC/D3 is much lower than in human brain microvessels (Eigenmann et al., 2013; Ohtsuki et al., 2013; Weksler et al., 2013), which may cause the low TEER values. VE-cadherin, a major member of the cadherin family in endothelial cells, is also required for BBB integrity (Dejana et al., 2008; Li et al., 2018). The absence of VE-cadherin resulted in faulty cell-to-cell junctions and disrupted distribution of ZO-1 (Sauteur et al., 2017; Tunggal et al., 2005). Strongly negative correlations between *P_app_* values of fluorescein or FITC-Dex and expression of claudin-5 or VE-cadherin further demonstrated the importance of claudin-5 and VE-cadherin in BBB integrity.

Neurons and astrocytes, as important components of the NVU, may be involved in the formation and maintenance of BBB function. Several reports showed that co-culture with astrocytes CC-2565 or SC-1810 enhanced the TEER of hCMEC/D3 cells (Hatherell et al., 2011), whereas co-culture of RBE4.B cells with primary rat neurons and astrocytes showed a lower permeability of [^3^H] sucrose than mono-culture of RBE4.B cells (Schiera Gabriella. et al., 2005). Similarly, co-cultured with hCMEC/D3 cells and 1321N1 (astrocytes) or 1321N1+SH-SY5Y cells possess higher TEER values and lower permeability to Lucifer yellow than hCMEC/D3 alone. Compared with double co-culture of hCMEC/D3 and 321N1 cells, the triple co-culture of hCMEC/D3, 1321N1, and SH-SY5Y cells showed higher TEER values (Barberio et al., 2022). Our study also demonstrated that co-culture with U251 and/or SH-SY5Y cells significantly lowered BBB permeability and upregulated VE-cadherin or claudin-5 expression in hCMEC/D3 cells.

Next, we focused on the molecular mechanisms by which neurons and possibly astrocytes upregulated the VE-cadherin and claudin-5 expression in BMECs. Co-culture with SH-SY5Y cells significantly upregulated claudin-5 and VE-cadherin expression in hCMEC/D3 cells. In the double co-culture with SH-SY5Y cells or triple co-culture BBB models, hCMEC/D3 cells were not in direct contact with SH-SY5Y cells, indicating that the interaction between SH-SY5Y and hCMEC/D3 cells depended on the release of some active compounds. It was consistent with the above deduction that the S-CM also markedly induced claudin-5 and VE-cadherin expression. Different from S-CM, U-CM mainly upregulated the VE-cadherin expression and just had a slight impact on claudin-5. In general, neurons but also astrocytes secrete some neurotrophic factors (Lonka-Nevalaita et al., 2010; Sweeney et al., 2019) that could contribute to the maintenance of structural stability of BBB. High levels of bFGF, GDNF, IGF-1, and TGF-*β* were detected in U-CM, S-CM, and US-CM. Further research showed that only GDNF induced VE-cadherin and claudin-5 expression in hCMEC/D3 cells in a concentration-dependent manner, and the anti-GDNF antibody attenuated claudin-5 and VE-cadherin expression induced by US-CM or GDNF. These results indicated that neurons upregulated claudin-5 and VE-cadherin expression through GDNF secretion. Levels of GDNF in S-CM were higher than those in U-CM, which seemed to partly explain why S-CM has a stronger promoting effect on claudin-5 than U-CM. The roles of GDNF in BBB maintenance and the regulation of claudin-5 and VE-cadherin expression were further confirmed using brain-specific *Gdnf* knockdown C57BL/6J mice. Consistent with our *in vitro* results, brain-specific *Gdnf* silencing greatly increased BBB penetration of fluorescein and FITC-Dex, accompanied by the downregulation of claudin-5 and VE-cadherin expression.

GDNF is mainly expressed in astrocytes and neurons (Lonka-Nevalaita et al., 2010; Pochon et al., 1997). In adult animals, GDNF is mainly secreted by striatal neurons rather than astrocytes and microglial cells (Hidalgo-Figueroa et al., 2012). The present study also shows that GDNF mRNA levels in SH-SY5Y cells were significantly higher than that in U251 cells. GDNF was also detected in conditioned medium from SH-SY5Y cells. All these results demonstrate that neurons may secrete GDNF.

Generally, GDNF activates several signal transduction pathways, such as the PI3K/AKT and MAPK signaling pathways (Fielder et al., 2018) by forming a heterohexameric complex with two GFR*α* molecules and RET receptors. It was also reported that GDNF improved BBB barrier function due to the activation of MAPK/ERK1 (Dong et al., 2018) and PI3K/AKT (Liu D. et al., 2022) signaling. Consistent with previous reports, we found that signaling inhibitors SPP-86, PP2, LY294002, and U0126 markedly attenuated US-CM- and GDNF-induced claudin-5 and VE-cadherin expression in hCMEC/D3 cells, inferring that GDNF promoted claudin-5 and VE-cadherin expression via activating both the PI3K/AKT and MAPK/ERK pathways.

Signal transduction pathways control gene expression by modifying the function of nuclear transcription factors. The nuclear accumulation of FOXO1 negatively regulates claudin-5 expression (Beard et al., 2020; Taddei et al., 2008). FOXO1 is an important target of the PI3K/AKT signaling axis (Zhang et al., 2011), and AKT-induced phosphorylation of FOXO1 results in cytoplasmic FOXO1 accumulation and decreases nuclear FOXO1 accumulation (Zhang et al., 2011). From these results, we inferred that GDNF-induced claudin-5 expression in hCMEC/D3 cells may be involved in the activation of PI3K/AKT/FOXO1 pathway. Similarly, both US-CM and GDNF increased cytoplasmic p-FOXO1 and decreased nuclear FOXO1 in hCMEC/D3 cells, which was reversed by LY29002 rather than U0126. Roles of FOXO1 in GDNF-induced claudin-5 were verified through silencing and overexpressing *FOXO1* in hCMEC/D3 cells. *FOXO1* silencing enhanced claudin-5 but not VE-cadherin expression, accompanied by a decline in total and nuclear FOXO1. In contrast, *FOXO1* overexpression significantly decreased claudin-5 expression. In hCMEC/D3 cells overexpressing *FOXO1*, GDNF lost its promotion effect on claudin-5 expression. It was noticed that U0126 attenuated the GDNF-induced upregulation of claudin-5 but had minimal impact on GDNF-mediated FOXO1 phosphorylation and the decline of nuclear FOXO1. In Sertoli cells, it was found that testosterone stimulated claudin-5 expression by activating the RAS/RAF/ERK/CREB pathway (Bulldan et al., 2016). The transcriptional regulation of claudin-5 by CREB was confirmed in bEnd.3 (mouse brain endothelial cell). CREB overexpression significantly increased both gene and protein expression of claudin 5. In contrast, depletion of CREB decreased claudin-5 expression in gene and protein levels (Li Y. et al., 2022). However, another report showed that in human lung microvascular endothelial cells, U0126 attenuated phosphorylation of ERK and lipopolysaccharide-stimulated claudin-5 damage, indicating activation of MAPK/ERK pathway impaired rather than promoted claudin-5 expression (Liu Y. et al., 2019). Thus, the real mechanisms that GDNF-induced activation of the RET/MAPK/ERK pathway promotes claudin-5 expression need further investigation.

ETS1 is a member of the ETS family that plays an important role in cell adhesion, migration, and blood vessel information. ETS1 binds to an ETS-binding site located in the proximal region of the VE-cadherin promoter, controlling VE-cadherin expression (Lelièvre et al., 2001). Several studies have demonstrated the role of ETS1 in the regulation of VE-cadherin expression. For example, IFN-*γ* and TNF-*α* impaired BBB integrity by decreasing ETS1-induced VE-cadherin expression (Luo et al., 2022). *ETS1* silencing reduced VE-cadherin expression in umbilical vein endothelial cells (Colas-Algora et al., 2020). In contrast, *ETS1* overexpression induced VE-cadherin expression in mouse brain capillary endothelial cells and fibroblasts (Lelièvre et al., 2000). As expected, *ETS1* silencing resulted in a decrease in the expression of VE-cadherin in hCMEC/D3 cells, but claudin-5 expression remained unaffected. Additionally, *ETS1* silencing removed the inductive effect of GDNF on VE-cadherin expression while unaffecting the upregulation of claudin-5 induced by GDNF. The activation of the PI3K/AKT (He et al., 2023; Hui et al., 2018) and MAPK/ERK (Watanabe et al., 2004) pathways promotes ETS1 expression. In our findings, US-CM and GDNF significantly increased total and nuclear ETS1 levels, which were eliminated by signaling inhibitors LY294002 and U0126. Both our results and previous research provide evidence that GDNF upregulates ETS1 expression via the activation of PI3K/AKT and MAPK/ERK signaling to promote VE-cadherin expression.

Claudin-5 expression is also regulated by VE-cadherin (Taddei et al., 2008). Differing from the previous reports, silencing VE-cadherin with siRNA only slightly affected basal and GDNF-induced claudin-5 expression. The discrepancies may come from different characteristics of the tested cells. Several reports have supported the above deduction. In retinal endothelial cells, hyperglycemia remarkably reduced claudin-5 expression (but not VE-cadherin) (Saker et al., 2014). However, in hCMEC/D3 cells, hypoglycemia significantly decreased claudin-5 expression but hyperglycemia increased VE-cadherin expression (Sajja et al., 2014).

The present study showed that characteristics of *in vitro* triple co-culture BBB model were superior to those of hCMEC/D3 mono-culture BBB model. The hCMEC/D3 mono-culture and triple co-culture BBB models were used to try to predict the *PS* values of 18 drugs by comparing them with their observations. The prediction success rate (14/18) of triple co-culture BBB model was greater than that of hCMEC/D3 mono-culture BBB model (7/18). However, poor predictions were observed for verapamil, amitriptyline, fluoxetine, and clozapine, which may partly be due to inaccuracies in their *f_u, brain_* values. These four drugs are high protein-binding drugs. Due to the methodological discordance and limitations of historic devices for these drugs, the *f_u_* ≤ 0.01 maybe with low confidence and accuracy (Bowman et al., 2018; Di et al., 2017). For example, *f_u, brain_* values of amitriptyline from different reports display large differences (0.002 (Sanchez-Dengra et al., 2021), 0.0071 (Weber et al., 2013)). The same large differences of *f_u, brain_* values were also found in fluoxetine (0.0023 (Maurer et al., 2005), 0.004 (Summerfield et al., 2007), 0.00094 (Liu X. et al., 2005)), and clozapine (0.0056 (Bhyrapuneni et al., 2018), 0.014 (Cremers et al., 2012), 0.0094 (Summerfield et al., 2007)).

In summary, our triple co-culture BBB model outperformed the mono-culture or double co-culture BBB models, mainly attributing to the fact that neurons and possibly astrocytes upregulated claudin-5 and VE-cadherin expression by secreting GDNF through activating PI3K/AKT and MAPK/ERK pathways. Additionally, the developed *in vitro* triple co-culture BBB model accurately predicted the *in vivo* BBB permeability of CNS drugs. This suggests the potential of our triple co-culture BBB model for utilization in CNS candidate screening during the drug development process (Figure 8).

**Figure 8.**
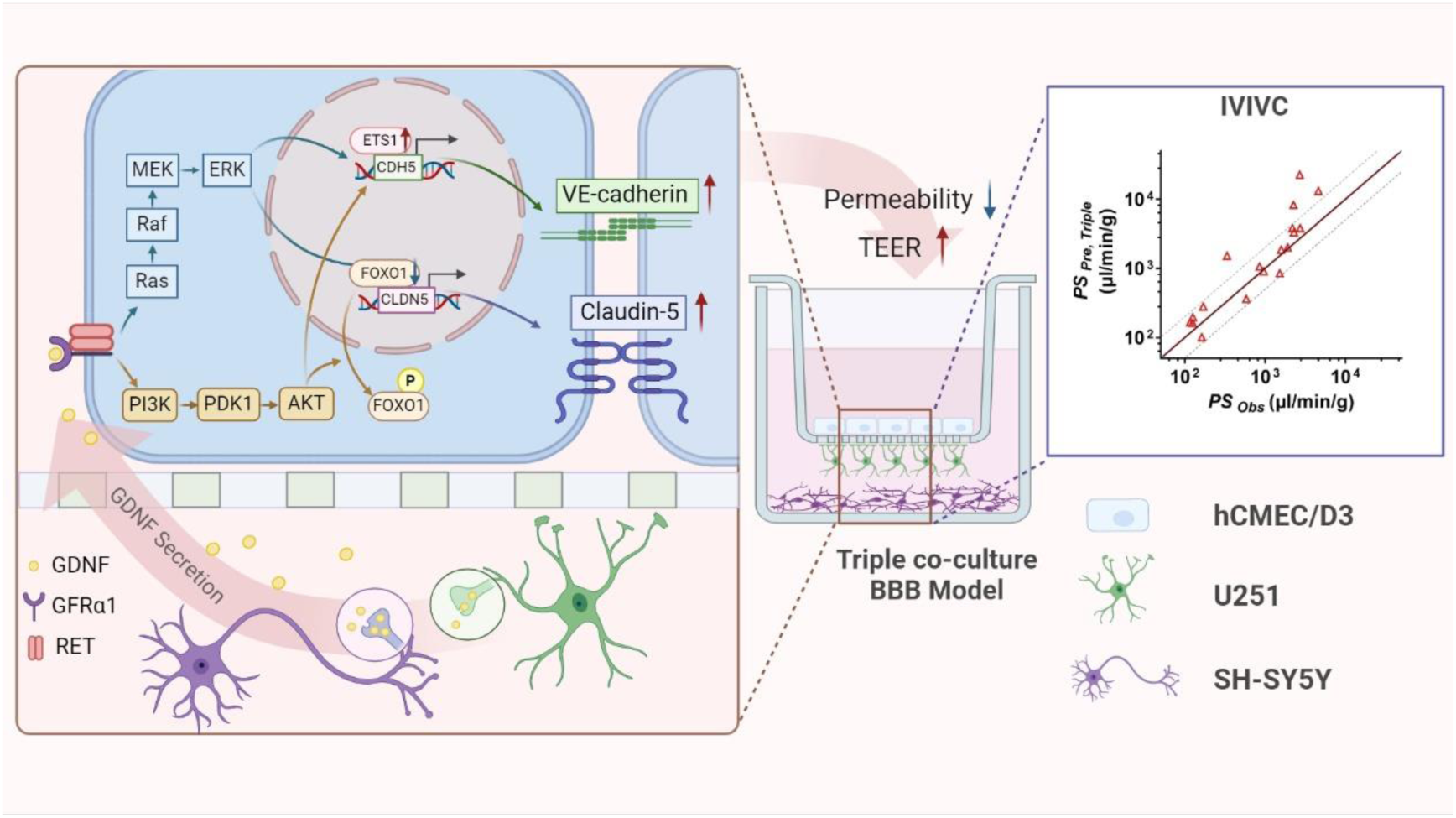
The mechanism of neurons and astrocytes induced the integrity of brain endothelial cells. Neurons but also astrocytes trigger the activation of PI3K/AKT and MAPK/ERK pathways in brain endothelial cells by GDNF secretion, which in turn regulates transcription factors of claudin-5 (FOXO1) and VE-cadherin (ETS1) to promote claudin-5 and VE-cadherin expression and leads to the enhancement of BBB integrity. Meanwhile, with the increase in barrier integrity, the *in vitro* BBB model also obtained a stronger *in vivo* correlation.

However, the study also has some limitations. In addition to neurons and astrocytes, other cells such as microglia, pericytes, and vascular smooth muscle cells, especially pericytes, may also affect BBB function. How pericytes affect BBB function and interaction among neurons, astrocytes, and pericytes needs further investigation.

## Materials and Methods

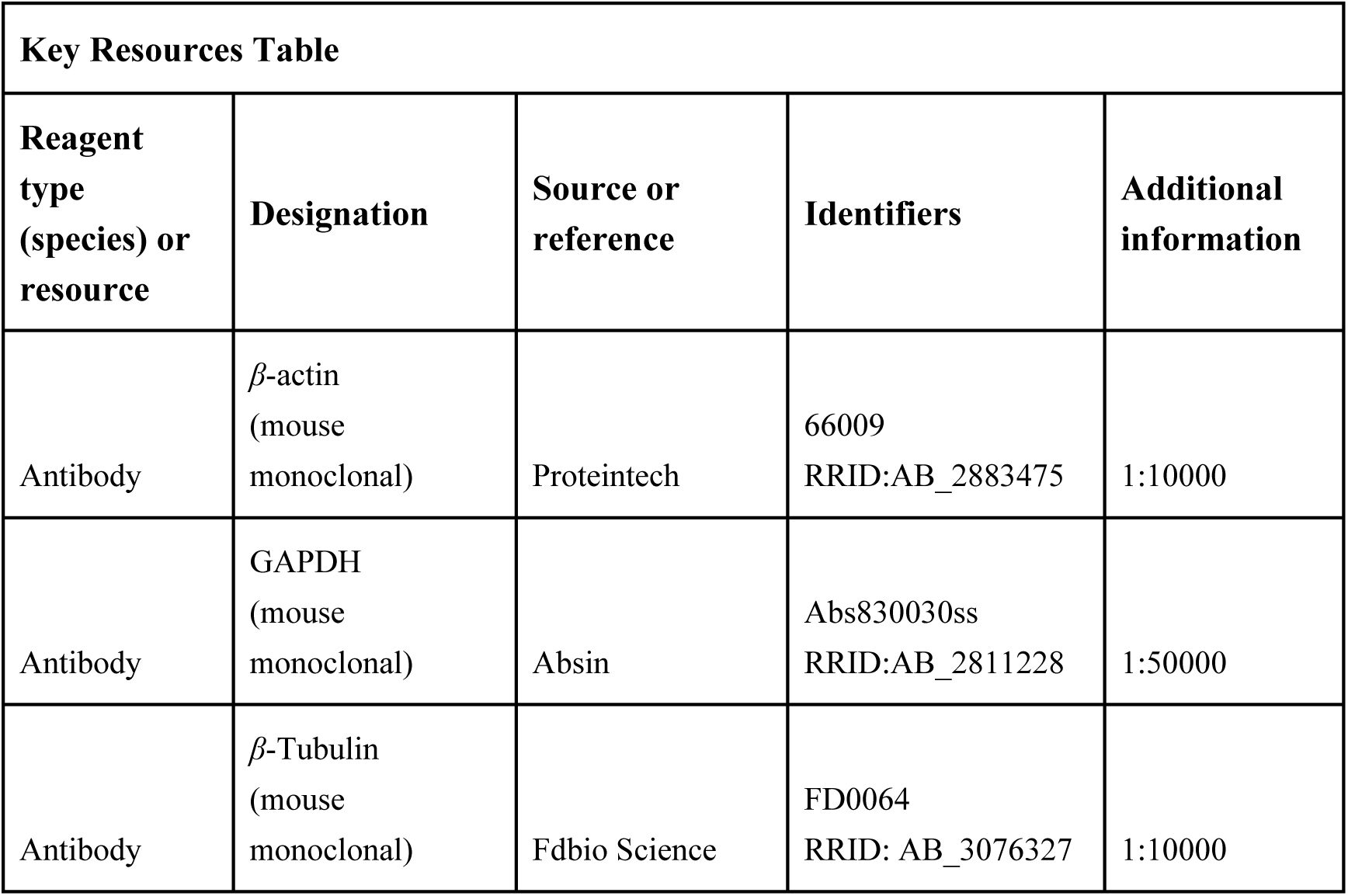
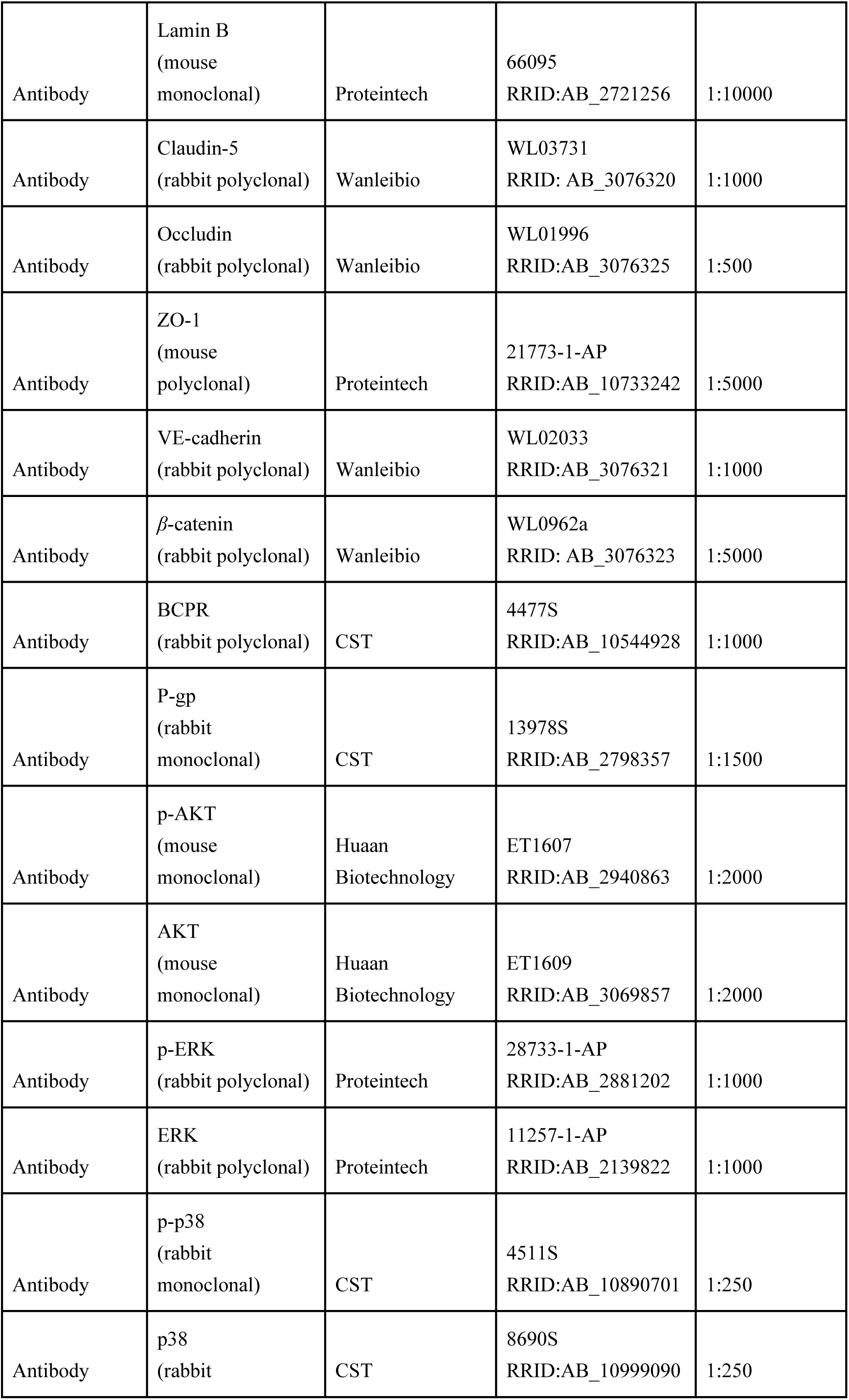
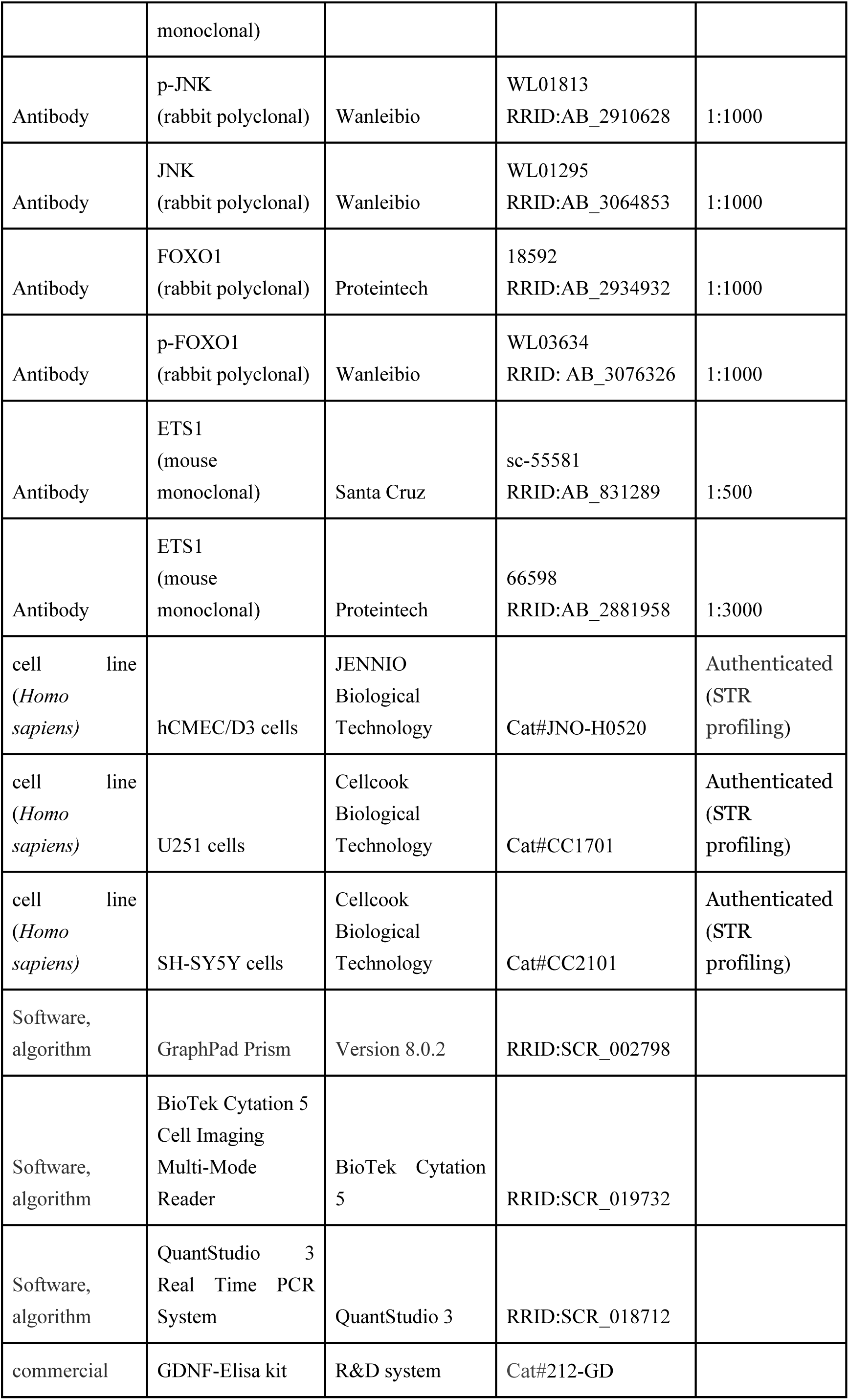
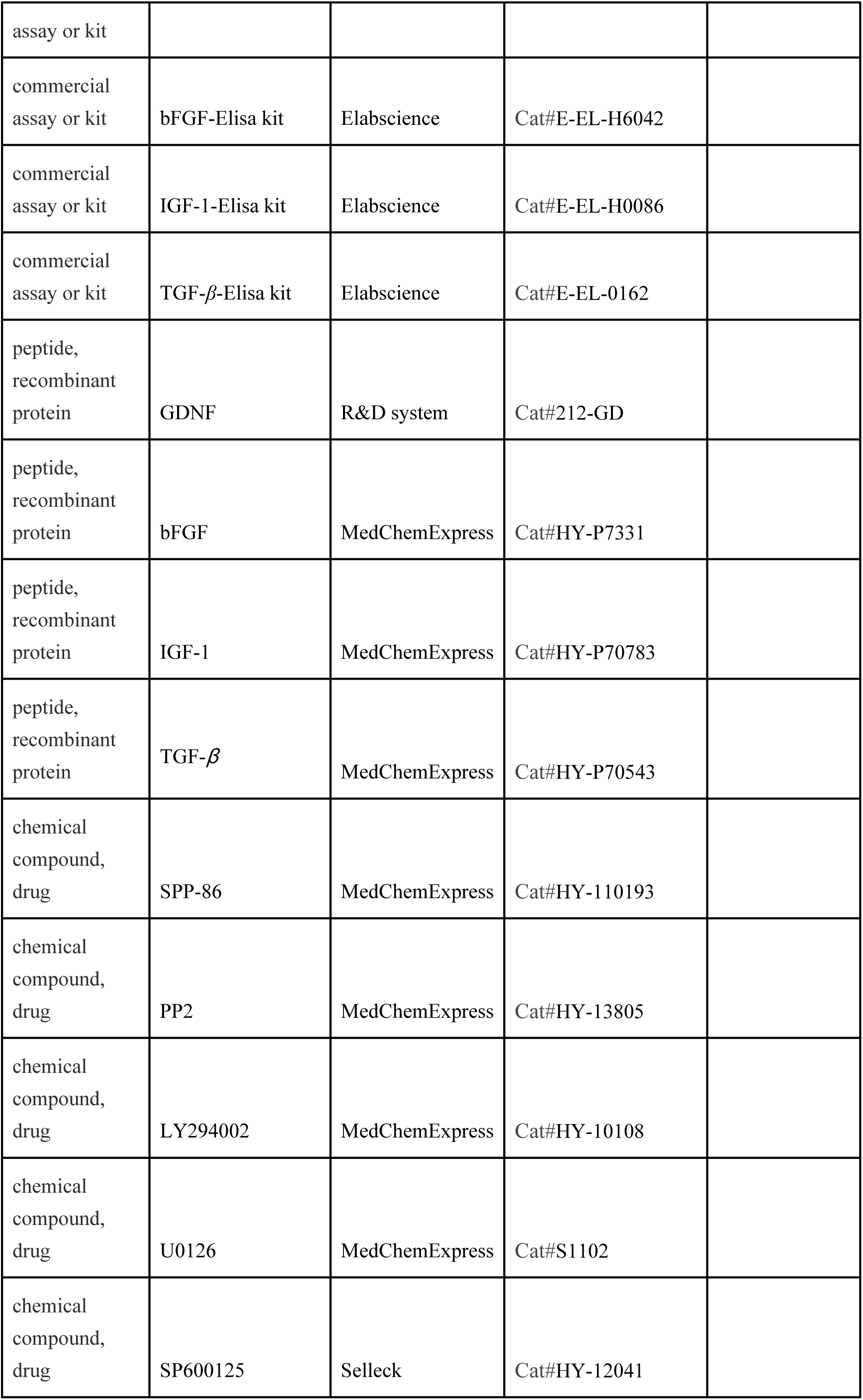
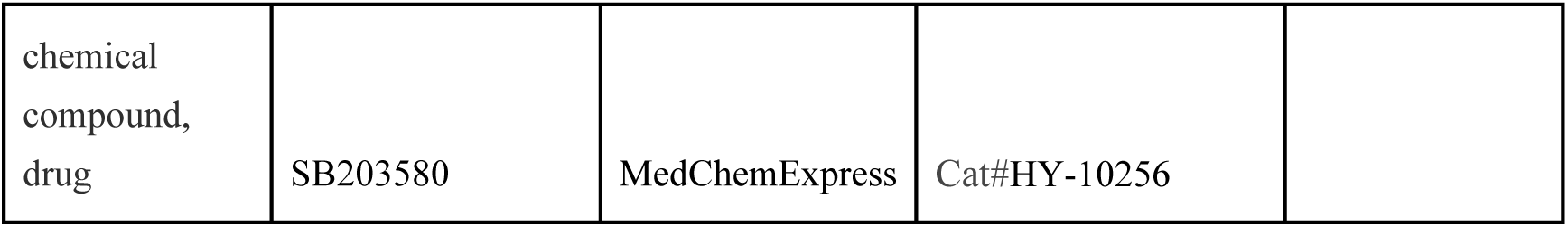

### Cell culture and viability assay

Rat brain microvascular endothelial cells were isolated from Sprague-Dawley rats (male, 7–10 days old, Sino-British Sippr/BKLaboratory Animal Ltd, Shanghai, China) as the described method (Bian-Sheng Ji et al., 2013; Ming-Shan L et al., 2013) and cultured in Dulbecco’s Modified Eagle Media (DMEM)/F12 (#12500-039, Gibco, Carlsbad, CA, USA) containing 10% fetal bovine serum (FBS) (#10100147C, Gibco, Carlsbad, CA, USA) and 62.5 µg/mL penicillin and 100 µg/mL streptomycin (SunShine Biotechnology Co., ltd. Nanjing, China). Then, hCMEC/D3, U251, and SH-SY5Y cells were cultured in DMEM/F12 containing 10% FBS, 62.5 µg/mL penicillin and 100 µg/mL streptomycin. Cell viability was assessed using a CCK-8 kit (Beyotime Biotechnology, Shanghai, China), and the results were expressed as the fold of control.

### Establishment of the triple co-culture model

Although hCMEC/D3 cells have poor barrier properties and low TEER compared to human physiological BBB, the use of human BMECs may be restricted by the acquisition of materials and ethical approval. Isolation and purification of primary BMECs are time-consuming and laborious. Moreover, culture conditions can alter transcriptional activity (Qi et al., 2023). All limit the establishment of BBB models based on primary human BMECs for high-throughput screening. Here, hCMEC/D3 cells were selected to establish an *in vitro* BBB model.

The establishment process of the triple co-culture model is illustrated in Figure 9. SH-SY5Y cells were seeded at a density of 4.5 × 10^4^ cells/cm^2^ in plates and differentiated with 10 μM retinoic acid (Sigma-Aldrich, St. Louis, MO, USA) for 72 h. The differentiated SH-SY5Y cells were cultured in the fresh DMEM/F12 medium containing 10% FBS. U251 cells were seeded at 2 × 10^4^ cells/cm^2^ on the bottom of Transwell inserts (PET, 0.4 µm pore size, SPL Life Sciences, Pocheon, Korea) coated with rat-tail collagen (Corning Inc., Corning, NY, USA). Next, the inserts were suspended in plate wells containing the culture medium after 5 h of incubation. After 24 h of incubation, hCMEC/D3 cells were seeded on the apical side of the inserts at 3 × 10^4^ cells/cm^2^ and cultured for another 48 h. Then, the inserts seeded with U251 and hCMEC/D3 cells were suspended in plates seeded with differentiated SH-SY5Y cells and co-cultured for another 6 days. The culture medium was replaced every 24 h. TEER was periodically measured using a Millicell-ERS (MERS00002) instrument (Millipore, Billerica MA, USA) to monitor cell confluence and development of tight junctions.

**Figure 9.**
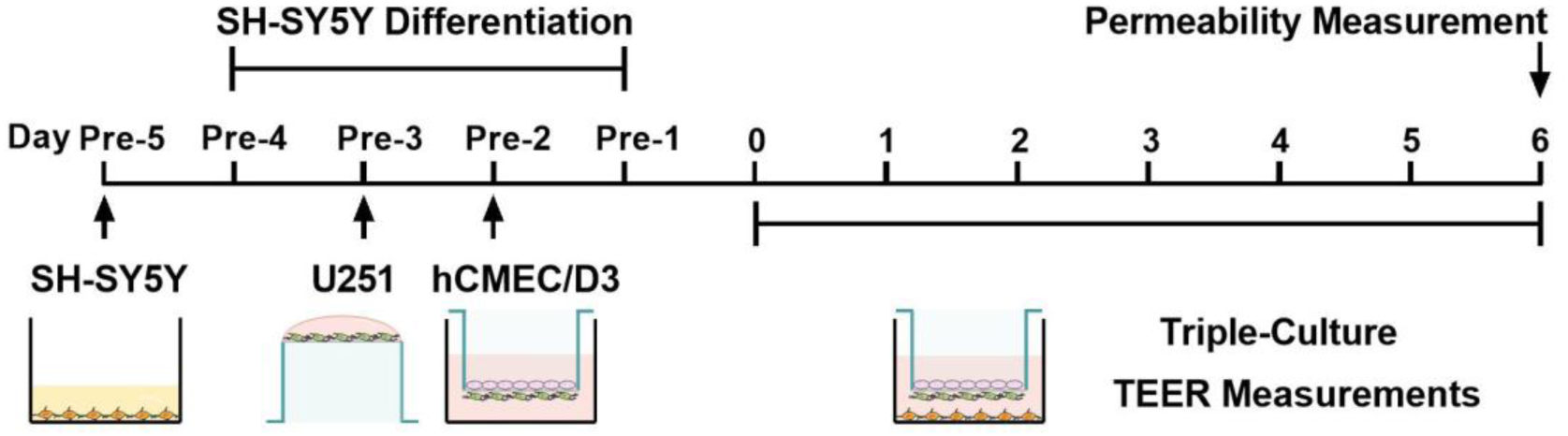
**Schematic diagram of the establishment process of the triple co-culture BBB model.**

### EdU incorporation assay

The cells were incubated with medium containing 10 μM Edu for 2 hours. Then cells were washed by PBS and harvested by 0.25% trypsin-ethylenediaminetetraacetic acid (#25200072, Gibco, Carlsbad, CA, USA). The EdU incorporation assay was measured using the BeyoClick™ EdU Cell Proliferation Kit (Beyotime Biotechnology, Shanghai, China) according to the manufacturer’s instructions. The samples were determined on the FACSCelesta flow cytometer (Becton, Dickins on and Company, USA, and data were analyzed by Flowjo 10.4 software.

### *In vitro* BBB permeability study

On day 7, the TEER values of BBB models showed a decreasing trend. Therefore, the subsequent experiments were all completed on day 6. The culture medium was removed from the apical and basolateral sides of the inserts and washed twice with preheated Hank’s balanced salt solution (HBSS). Fresh HBSS was then added to both the apical and basolateral chambers. After 15 min of preincubation, HBSS in the apical and basolateral chambers was replaced with HBSS containing FITC-dextran 3–5 kDa (FITC-Dex) (Sigma-Aldrich, St. Louis, MO, USA), fluorescein sodium (Sigma-Aldrich, St. Louis, MO, USA), or other tested agents and blank HBSS, respectively. Next, 200 μL aliquots were collected from the basolateral chamber after 30 min of incubation at 37 °C. The concentrations of the tested agents in the basolateral chamber were measured.

The apparent permeability coefficient (*P_app_*, cm/s) values of the tested agents across the *in vitro* BBB model were calculated using the equation (Tavelin et al., 2002):

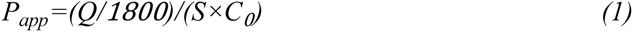

where *S* is the surface area of the insert membrane (0.33 cm^2^ for 6.5 mm inserts, 4.46 cm^2^ for 24 mm inserts), *Q* is the transported amount of the tested agents transported from the donor chamber to the receiver chamber for 30 min (1800 s), and *C_0_* is the initial concentration of the tested agents in the donor chamber.

### The quantification methods of prazosin, verapamil, lamotrigine

Prazosin, verapamil, and lamotrigine (Aladdin, Shanghai, China) were analyzed by high-performance liquid chromatography (Shimadzu, Kyoto, Japan) with YMC-Triart C18 column (5 μm, 150 × 4.6 mm, YMC America Inc., Allentown, PA, USA). Prazosin and verapamil were detected using the RF-20A fluorescence detector. Lamotrigine was detected using the SPD-20A ultraviolet detector. Samples were centrifugated at 12000 rpm for 10 mins, then 150 μL supernatant was taken and used for analysis. The run temperature was set at 40 ℃, the injection volume was 20 μL and the flow rate was 1 mL/min. Initial concentrations in donor chamber and other chromatographic conditions of drugs are summarized in Table 2.

**Table 2.**
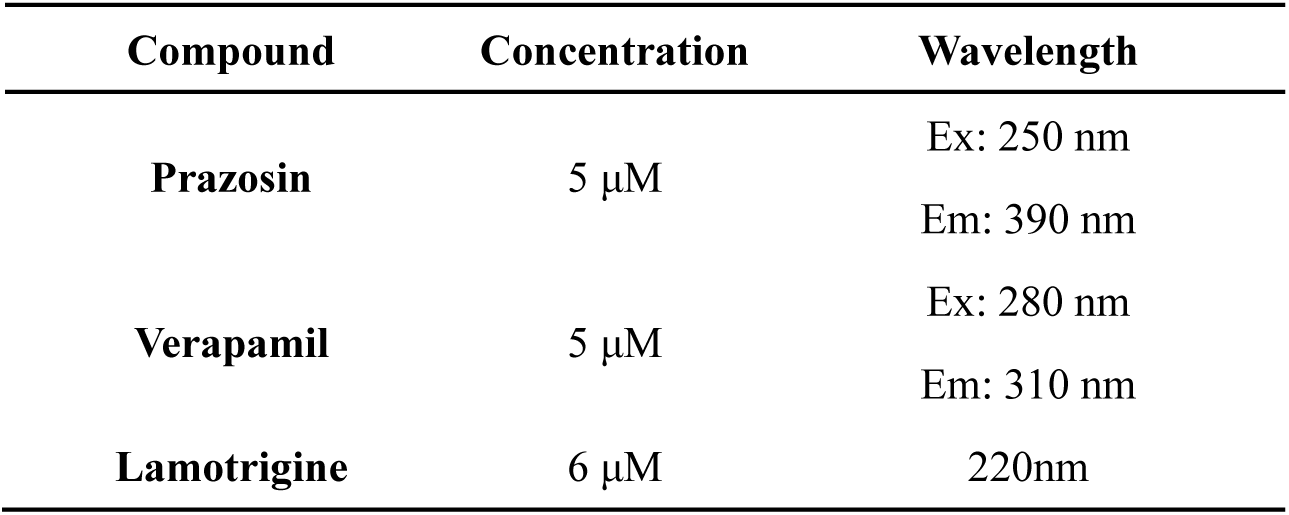
Initial concentrations in donor chamber and chromatographic conditions of prazosin, verapamil, and lamotrigine.

### The quantification methods of clozapine, venlafaxine, bupropion, amantadine, carbamazepine, fluoxetine, amitriptyline, gabapentin, midazolam, risperidone, olanzapine, mirtazapine, metoclopramide, doxepin, donepezil

All compounds were purchased from Aladdin (Shanghai, China). Except for prazosin, verapamil, and lamotrigine, the other compounds were analyzed by using liquid chromatography-mass spectrometry (Shimadzu, Kyoto, Japan) with YMC-Triart C18 column (5 μm, 150 × 2.0 mm, YMC America Inc., Allentown, PA, USA). Each sample was mixed with 10 µL internal standard. Then 1 mL extraction was added to each sample. The samples were vortex vibrated on the oscillator for 10 min, and then centrifuged at 4 ℃ and 12000 rpm for 10 mins. The supernatant solvent was evaporated with nitrogen flow, then redissolved with 100 µL 40 % (v/v) acetonitrile, and centrifuged at 4 ℃ and 15000 rpm for 10 min. The supernatants were injected into LC-MS for analysis. The injection volume of each sample was 5 µL. The mass charge ratio, extraction, and initial concentrations in donor chamber of drugs were summarized in Table 3.

**Table 3.** The summary of mass charge ratio, extraction, initial concentrations in donor chamber.

| Compound | Concentration | Mass charge ratio<br>[M+H]<sup>+</sup> | Extraction |
| --- | --- | --- | --- |
| Amantadine | 3 μM | 181 | Water-saturated N-butanol |
| Amitriptyline | 1.5 μM | 278 | Ethyl acetate |
| Bupropion | 3 μM | 240 | Ethyl acetate |
| Carbamazepine | 3 μM | 237 | Ethyl acetate |
| Clozapine | 4 μM | 327 | Ethyl acetate |
| Donepezil | 3 μM | 380 | Methyl tert-butyl ether |
| Doxepin | 4 μM | 317 | Methyl tert-butyl ether |
| Fluoxetine | 3 μM | 310 | Ethyl acetate |
| Gabapentin | 10 μM | 172 | Ethyl acetate |
| Metoclopramide | 4 μM | 301 | Ethyl acetate |
| Midazolam | 3 μM | 327 | Ethyl acetate |
| Mirtazapine | 3 μM | 266 | Methyl tert-butyl ether |
| Olanzapine | 3 μM | 313 | Methyl tert-butyl ether |
| Risperidone | 4 μM | 427 | Methyl tert-butyl ether |
| Venlafaxine | 10 μM | 278 | Ethyl acetate |

### Cell density analysis

On day 6 of co-culture, hCMEC/D3 cells were fixed with 4% paraformaldehyde for 15 min and washed with phosphate buffered saline (PBS) for three times. Next, the fixed cells were blocked with 5% goat serum for 2 h and washed with PBS for four times. The blocked cells were incubated with DAPI (Invitrogen, Carlsbad, CA, USA) and washed with PBS for four times. Cell numbers were counted using Cytation5 (BioTek, Winooski, VT, USA).

### Reverse transcription and Quantitative real-time PCR (qPCR)

Total RNA of cells was extracted using RNAiso Plus reagent (Takara Bio Inc. Otsu, Shiga, Japan) and reverse transcribed using HiScript III RT SuperMix (Vazyme, Shanghai, China) as the described method (Yang et al., 2023). Paired primers were synthesized by Tsingke Biotech Co., Ltd (Beijing, China), and their sequences were listed in Table 4. The SYBR Master Mix was purchased from Yeasen (Shanghai, China). Then, qPCR was performed on the Applied Biosystems QuantStudio 3 real-time PCR system (Thermo Fisher Scientific, Waltham, MA, USA). The mRNA levels of related genes were normalized to *ACTB* or *GAPDH* using the comparative cycle threshold method.

**Table 4.**
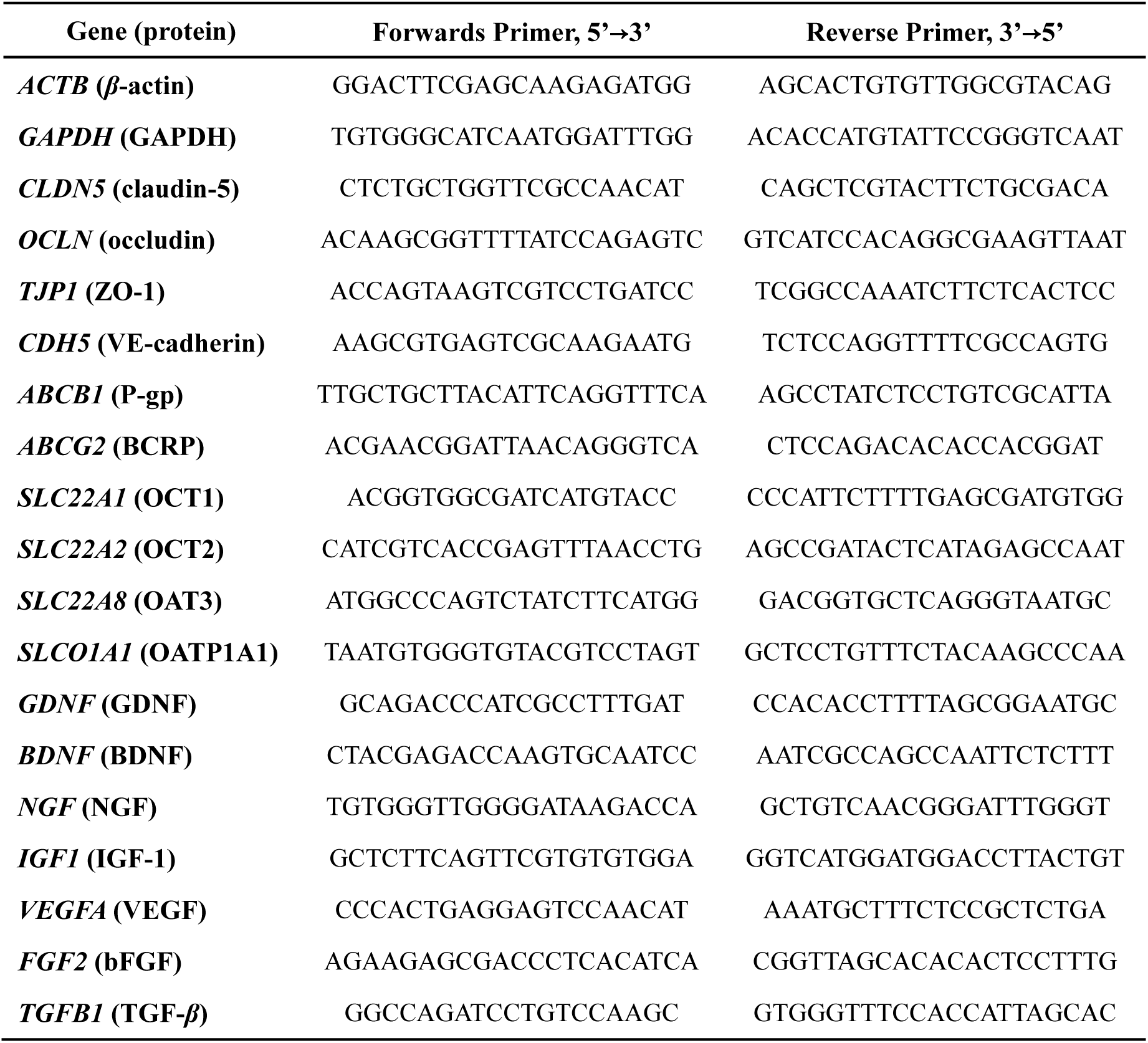
Primer sequences for qPCR for indicted genes.

### Western blotting analysis

Whole-cell and tissue lysates, nucleoprotein, and cytoplasmic protein were prepared using RIPA Lysis Buffer (Beyotime, Shanghai, China) as the described method (Wu et al., 2021). Proteins were separated through sodium dodecyl sulfate-polyacrylamide gel electrophoresis and transferred onto nitrocellulose or polyvinylidene difluoride membranes. The membranes were blocked with 5% skim milk and incubated with corresponding primary antibodies at 4 ℃ overnight. After being washed with Tris-buffered saline Tween buffer, the membranes were incubated with secondary antibodies (Cell Signaling Technology, MA, USA) at 1:3000 dilution: Anti-mouse IgG, HRP-linked Antibody (#7076), Anti-rabbit IgG, HRP-linked Antibody (#7074). Protein levels were visualized using a highly sensitive ECL western blotting substrate and a gel imaging system (Tanon Science & Technology, Shanghai, China).

### Preparation of conditioned medium

Conditioned medium (CM) of U251 cells (U-CM), SH-SY5Y cells (S-CM), or co-culture of U251 and SH-SY5Y cells (US-CM) were prepared. U251 cells were seeded at the top of the insert membrane and suspended on 6-well plates seeded with differentiated SH-SY5Y cells to co-culture U251 cells with SH-SY5Y cells. The medium was collected every 24 h. The CMs were subsequently used for hCMEC/D3 cell culture after filtrating with 0.2 μm filters for 144 h. The levels of glia-derived neurotrophic factor (GDNF), basic fibroblast growth factor (bFGF), insulin-like growth factor-1 (IGF-1), and transforming growth factor-*β* (TGF-*β*) in CMs were measured using corresponding ELISA kits according to the manufacturers’ instructions.

### Neutralization of GDNF with Anti-GDNF Antibody

Exogenous and endogenous GDNF in the medium was neutralized with anti-GDNF antibody (#AF-212-NA, R&D system, Minneapolis, MN, USA). 0.25, 0.5, 1.0 μg/mL anti-GDNF antibody was added into the US-CM or medium containing 200 pg/mL GDNF, and then the medium was preincubated at 4 ℃ for 1 h. The hCMEC/D3 cells were incubated with 200 pg/mL GDNF or US-CM containing anti-GDNF antibody or not for 6 days, and then cell lysate was collected for western blot. The medium was replaced every 24 h.

### Transfection of hCMEC/D3

Here, hCMEC/D3 cells were plated in the plates or culture dishes at 6 × 10^4^ cells/cm^2^ and transfected with 10 nM of negative control or human *FOXO1* and *ETS1* small interfering RNA (siRNA) (Tsingke Biotechnology, Beijing, China) for 12 h using Lipofectamine™ 3000 (Invitrogen, Carlsbad, CA, USA) reagent according to the manufacturer’s instructions. Cells were then incubated with a medium containing GDNF for 72 h. The siRNA sequences of human *FOXO1* and *ETS1* were summarized in Table 5.

**Table 5.**
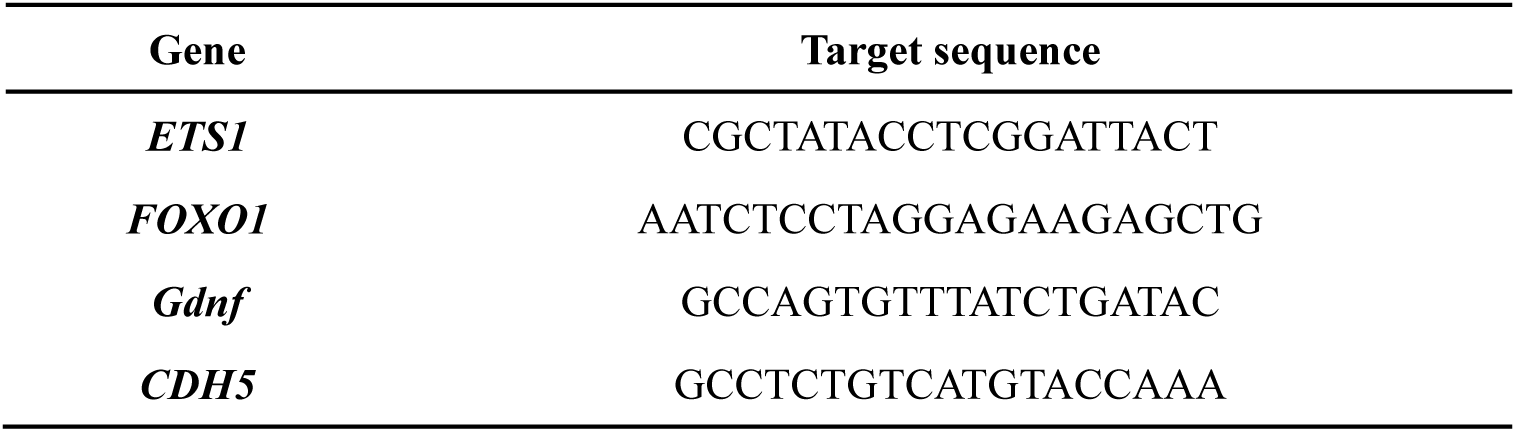
The target sequences for siRNA or shRNA.

### FOXO1 Overexpression by plasmids

The plasmids encoding FOXO1 (EX-Z7404-M02) were constructed by GeneCopoeia (Rockville, MD, USA). The hCMEC/D3 cells were plated in plates or dishes at 6 × 10^4^ cells/cm^2^. They were subsequently transfected with 1 µg of negative control or plasmids encoding FOXO1 6 h using Lipofectamine™ 3000 reagent according to the manufacturer’s instructions. Transfected cells were then incubated with the medium containing GDNF for 72 h.

### Animals

C57BL/6J mice (male, 4–5 weeks old, 16–18 g, 12 mice) were obtained from Sino-British Sippr/BKLaboratory Animal Ltd (Shanghai, China). Mice were maintained in groups under standard conditions with free access to food and water. Animal studies were performed in accordance with the Guide for the Care and Use of Laboratory Animals (National Institutes of Health) and approved by the Animal Ethics Committee of China Pharmaceutical University (Approval Number: 202307003).

### Brain-specific *Gdnf* knockdown and evaluation of BBB permeability

Mice were randomly divided into control (shNC) and *Gdnf* silencing (sh*Gdnf*) groups (6 mice each group). The shRNA sequence of mice *Gdnf* was listed in Table 5. The 2 × 10^9^ viral genome each of pAAV-U6-shRNA (NC2)-CMV-EGFP or pAAV- U6-shRNA (*Gdnf*)-CMV-EGFP (OBio Technology, Beijing, China) were injected into the bilateral lateral ventricle area (relative to the bregma: anterior-posterior -0.3 mm; medial-lateral ±1.0 mm; dorsal-ventral -3.0 mm) through intracerebroventricular (*i.c.v*) infusion. Three weeks following *i.c.v* injection, BBB permeability and expression of corresponding targeted proteins were measured in the mice.

A mixture of FITC-Dex (50 mg/kg) and fluorescein sodium (10 mg/kg) was intravenously administered to experimental mice. Thirty minutes after the injection, the mice were euthanized under isoflurane anesthesia, and brain tissue and plasma samples were obtained quickly. The concentrations of FITC-Dex and fluorescein in the plasma and brain were measured as previously described (Li P. et al., 2022; Zhou Y. et al., 2019). No blinding was performed in animal studies.

### The prediction of drug permeability across BBB using the developed in vitro BBB model

The *P_app_* values of 18 drugs – prazosin, verapamil, lamotrigine, clozapine, venlafaxine, bupropion, amantadine, carbamazepine, fluoxetine, amitriptyline, gabapentin, midazolam, risperidone, olanzapine, mirtazapine, metoclopramide, doxepin, and donepezil – across the hCMEC/D3 cells mono-culture and triple co-culture models were measured. The predicted *in vivo* permeability-surface area product (*PS*, μL/min/g brain) values across BBB were calculated using the following equation:

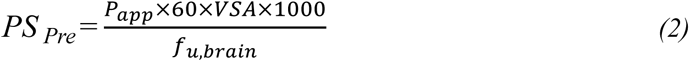

where VSA is the luminal area of the vascular space of brain, which was set to 150 cm^2^/g (Fenstermacher et al., 1988), and *f_u, brain_* is the unbound fraction of brain. The published *in vivo* brain permeability values were unified to observed *PS* (*PS _Obs_*) by multiplying by VSA equal to 150 cm^2^/g. If *PS _Pre_* values were within 0.5–2.0 folds of observations, the prediction was considered successful.

### Statistical analyses

All results are presented as mean ± SEM. The average of technical replicates generated a single independent value that contributes to the n value used for comparative statistical analysis. The data were assessed for Gaussian distributions using Shapiro-Wilk test. Brown-Forsythe test was employed to evaluate the homogeneity of variance between groups. For comparisons between two groups, statistical significance was determined by unpaired 2-tailed t-test. The acquired data with significant variation were tested using unpaired t-test with Welch’s correction, and non-Gaussian distributed data were tested using Mann-Whitney test. For multiple group comparisons, one-way ANOVA followed by Fisher’s LSD test was used to determine statistical significance. The acquired data with significant variation were tested using Welch’s ANOVA test, and non-Gaussian distributed data were tested using Kruskal-Wallis test. *P* < 0.05 was considered statistically significant. The simple linear regression analysis was used to examine the presence of a linear relationship between two variables. Data were analyzed using GraphPad Prism software version 8.0.2 (GraphPad Software, La Jolla, CA, USA).

## Acknowledgments

The authors would like to give special thanks to the support of the China Pharmaceutical University Pharmaceutical Animal Laboratory Center.

## Funding information

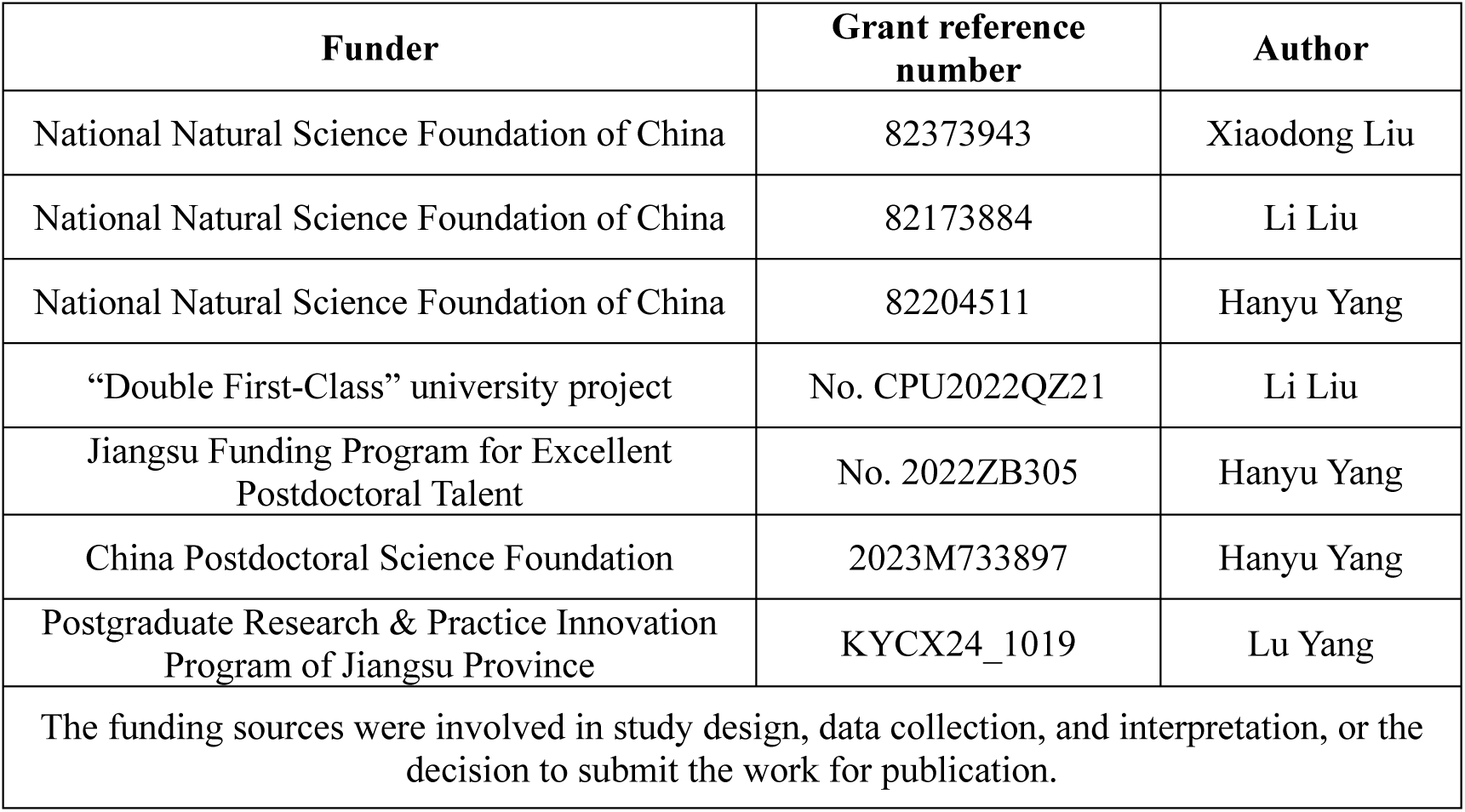

## Author Contributions

**Lu Yang:** Conceptualization, Methodology, Investigation, Data curation, and Writing–original draft preparation, Funding acquisition; **Zijin Lin:** Data curation, Formal analysis; **Ruijing Mu**: Visualisation, Software; **Wenhan Wu**: Formal analysis; **Hao Zhi**: Data curation; **Xiaodong Liu**: Supervision, Conceptualization, Project administration, Funding acquisition**; Hanyu Yang**: Supervision, Conceptualization, Funding acquisition**; Li Liu**: Supervision, Conceptualization, Project administration, Funding acquisition, Resources. All authors reviewed and approved the final version of this manuscript.

## Data availability

All data are available in the manuscript and supporting files; source data files for western blots have been provided for all figures.

---

**Figure 1—figure supplement 1:**
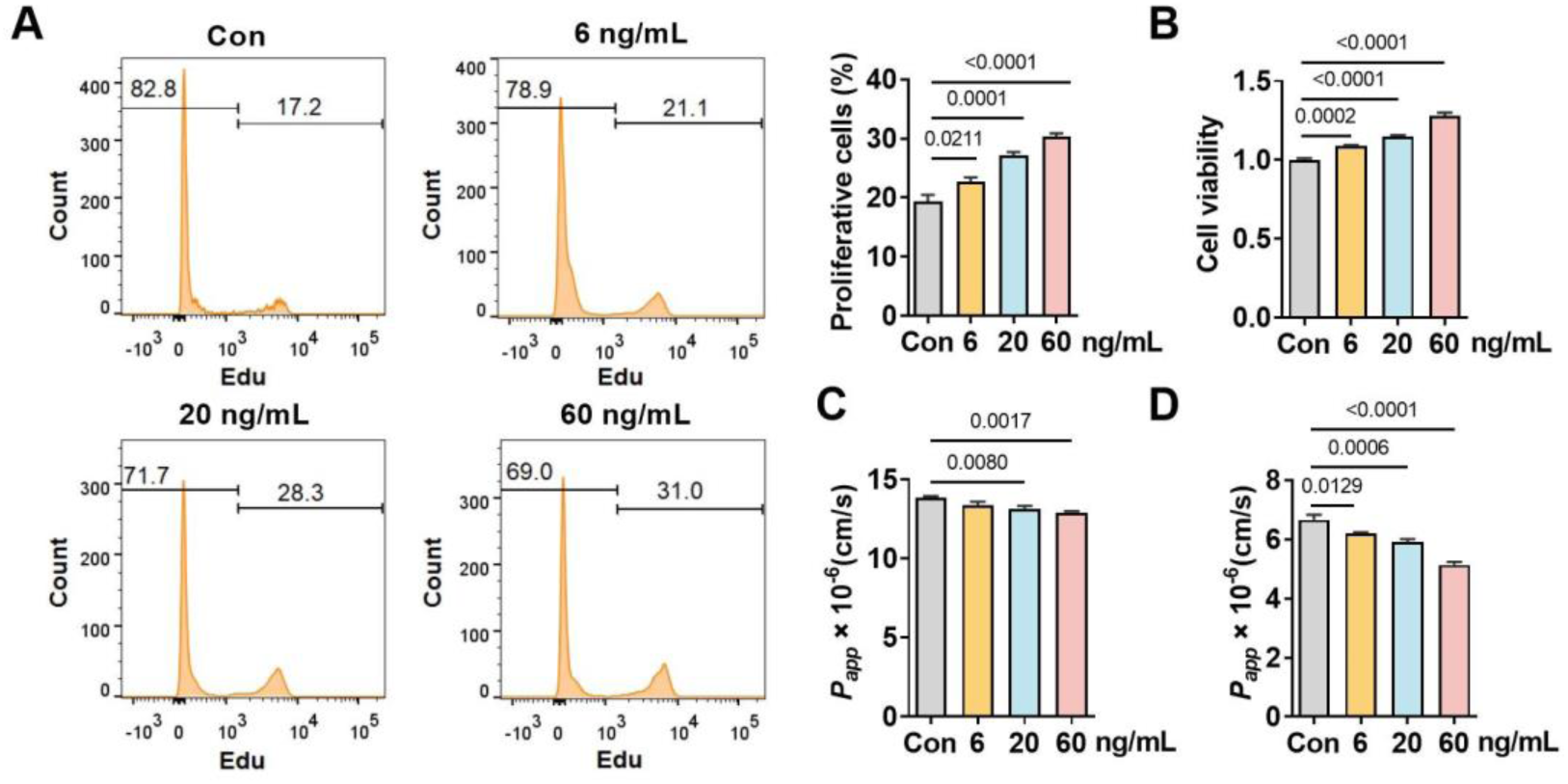
The induced proliferation of hCMEC/D3 cells by bFGF slightly reduced the permeability of cell layers. EdU incorporation (A), cell viability (B), and apparent permeability coefficient (*P_app_*, × 10^-6^ cm/s) of fluorescein (C) or FITC-Dextran 3–5 kDa (D) of hCMEC/D3 cells treated with bFGF (6, 20, 60 ng/mL) for 6 days. The above data are shown as the mean ± SEM. Four biological replicates per group. One technical replicate for each biological replicate. Statistical significance was determined using one-way ANOVA test followed by Fisher’s LSD test or Welch’s ANOVA test.

**Figure 4—figure supplement 1:**
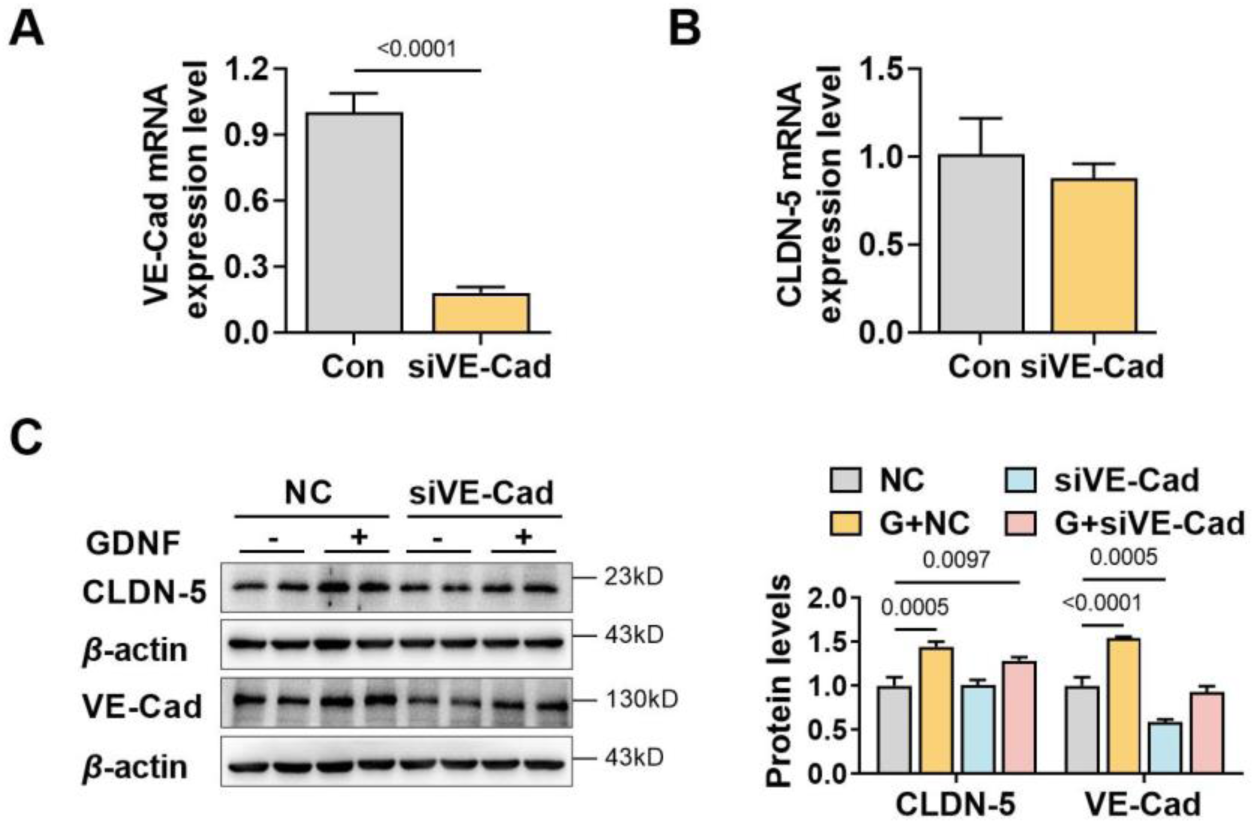
The contribution of VE-Cadherin on the GDNF-induced claudin-5 expression. Effects of the VE-Cadherin siRNA (siVE-Cad) on mRNA expression of VE-cadherin (A) and claudin-5 (B). Effects of siVE-Cad and GDNF on claudin-5 and VE-cadherin protein expression (C). NC: negative control plasmids. The above data are shown as the mean ± SEM. Four biological replicates per group. Two technical replicates for A and B and one technical replicate for C. Statistical significance was determined using unpaired Student’s t-test or one-way ANOVA test followed by Fisher’s LSD test.

**Figure 1-Source data1**

The western blot raw images in Figure 1

**Figure 1-Source data2**

The labeled western blot images in Figure 1

**Figure 1-Source data3**

Excel file containing summary data and data analysis of Figure 1

**Figure 2-Source data1**

The western blot raw images in Figure 2

**Figure 2-Source data2**

The labeled western blot images in Figure 2

**Figure 2-Source data3**

Excel file containing summary data and data analysis of Figure 2

**Figure 3-Source data1**

The western blot raw images in Figure 3

**Figure 3-Source data2**

The labeled western blot images in Figure 3

**Figure 3-Source data3**

Excel file containing summary data and data analysis of Figure 3

**Figure 4-Source data1**

The western blot raw images in Figure 4

**Figure 4-Source data2**

The labeled western blot images in Figure 4

**Figure 4-Source data3**

Excel file containing summary data and data analysis of Figure 4

**Figure 5-Source data1**

The western blot raw images in Figure 5

**Figure 5-Source data2**

The labeled western blot images in Figure 5

**Figure 5-Source data3**

Excel file containing summary data and data analysis of Figure 5

**Figure 6-Source data1**

The western blot raw images in Figure 6

**Figure 6-Source data2**

The labeled western blot images in Figure 6

**Figure 6-Source data3**

Excel file containing summary data and data analysis of Figure 6

**Table 1-Source data1**

The apparent permeability coefficients of 18 tested drugs from mono or triple culture BBB model

## References

Abbott N J. (2013). Blood-brain barrier structure and function and the challenges for CNS drug delivery. J Inherit Metab Dis, 36(3), 437–449. 10.1007/s10545-013-9608-0

Arvanitis C D, Ferraro G B, Jain R K. (2020). The blood-brain barrier and blood-tumour barrier in brain tumours and metastases. Nat Rev Cancer, 20(1), 26–41. 10.1038/s41568-019-0205-x

Asada S, Daitoku H, Matsuzaki H, Saito T, Sudo T, Mukai H, Iwashita S, Kako K, Kishi T, Kasuya Y, Fukamizu A. (2007). Mitogen-activated protein kinases, Erk and p38, phosphorylate and regulate Foxo1. Cell Signal, *19*(3), 519-527. 10.1016/j.cellsig.2006.08.015

Avdeef A, Sun N. (2010). A New In Situ Brain Perfusion Flow Correction Method for Lipophilic Drugs Based on the pH-Dependent Crone-Renkin Equation. Pharmaceutical Research, 28(3), 517–530. 10.1007/s11095-010-0298-0

Banks W A. (2016). From blood-brain barrier to blood-brain interface: new opportunities for CNS drug delivery. Nat Rev Drug Discov, 15(4), 275–292. 10.1038/nrd.2015.21

Barberio C, Withers A, Mishra Y, Couraud P-O, Romero I A, Weksler B, Owens R M. (2022). A human-derived neurovascular unit in vitro model to study the effects of cellular cross-talk and soluble factors on barrier integrity. Frontiers in Cellular Neuroscience, 16. 10.3389/fncel.2022.1065193

Beard R S, Jr., Hoettels B A, Meegan J E, Wertz T S, Cha B J, Yang X, Oxford J T, Wu M H, Yuan S Y. (2020). AKT2 maintains brain endothelial claudin-5 expression and selective activation of IR/AKT2/FOXO1-signaling reverses barrier dysfunction. J Cereb Blood Flow Metab, 40(2), 374–391. 10.1177/0271678X18817512

Bhallamudi S, Roos B B, Teske J J, Wicher S A, McConico A, C M P, Sathish V, Prakash Y S. (2021). Glial-derived neurotrophic factor in human airway smooth muscle. J Cell Physiol, 236(12), 8184–8196. 10.1002/jcp.30489

Bhattacharya C, Masters A R, Bach C, Stratford R E, Jr. (2021). Population model analysis of chiral inversion and degradation of bupropion enantiomers, and application to enantiomer specific fraction unbound determination in rat plasma and brain. J Pharm Biomed Anal, 195, 113872. 10.1016/j.jpba.2020.113872

Bhyrapuneni G, Thentu J B, Palacharla V R C, Muddana N, Aleti R R, Ajjala D R, Nirogi R. (2018). A definite measure of occupancy exposures, seeking with non-radiolabeled in vivo 5-HT2A receptor occupancy and in vitro free fractions. *J Recept Signal Transduct Res*, *38*(4), 359-366. 10.1080/10799893.2018.1531888

Bian-Sheng Ji, Juan Cen, Ling He, Meng Liu, Yan-Qing Liu, Liu L. (2013). Modulation of P-glycoprotein in rat brain microvessel endothelial cells under oxygen glucose deprivation. J Pharm Pharmacol, 65(10), 1508–1517. 10.1111/jphp.12122

Biswas S, Cottarelli A, Agalliu D. (2020). Neuronal and glial regulation of CNS angiogenesis and barriergenesis. Development, 147(9). 10.1242/dev.182279

Bowman C M, Benet L Z. (2018). An examination of protein binding and protein-facilitated uptake relating to in vitro-in vivo extrapolation. Eur J Pharm Sci, 123, 502–514. 10.1016/j.ejps.2018.08.008

Bulldan A, Dietze R, Shihan M, Scheiner-Bobis G. (2016). Non-classical testosterone signaling mediated through ZIP9 stimulates claudin expression and tight junction formation in Sertoli cells. Cellular Signalling, 28(8), 1075–1085. 10.1016/j.cellsig.2016.04.015

Clasadonte J, Prevot V. (2017). The special relationship: glia–neuron interactions in the neuroendocrine hypothalamus. Nature Reviews Endocrinology, 14(1), 25–44. 10.1038/nrendo.2017.124

Colas-Algora N, Garcia-Weber D, Cacho-Navas C, Barroso S, Caballero A, Ribas C, Correas I, Millan J. (2020). Compensatory increase of VE-cadherin expression through ETS1 regulates endothelial barrier function in response to TNFalpha. Cell Mol Life Sci, 77(11), 2125–2140. 10.1007/s00018-019-03260-9

Cremers T I, Flik G, Hofland C, Stratford R E, Jr. (2012). Microdialysis evaluation of clozapine and N-desmethylclozapine pharmacokinetics in rat brain. Drug Metab Dispos, 40(10), 1909–1916. 10.1124/dmd.112.045682

Dejana E, Orsenigo F, Lampugnani M G. (2008). The role of adherens junctions and VE-cadherin in the control of vascular permeability. J Cell Sci, 121(Pt 13), 2115–2122. 10.1242/jcs.017897

Di L, Umland J P, Chang G, Huang Y, Lin Z, Scott D O, Troutman M D, Liston T E. (2011). Species independence in brain tissue binding using brain homogenates. Drug Metab Dispos, 39(7), 1270–1277. 10.1124/dmd.111.038778

Di L, Breen C, Chambers R, Eckley S T, Fricke R, Ghosh A, Harradine P, Kalvass J C, Ho S, Lee C A, Marathe P, Perkins E J, Qian M, Tse S, Yan Z, Zamek-Gliszczynski M J. (2017). Industry Perspective on Contemporary Protein-Binding Methodologies: Considerations for Regulatory Drug-Drug Interaction and Related Guidelines on Highly Bound Drugs. J Pharm Sci, 106(12), 3442–3452. 10.1016/j.xphs.2017.09.005

Di Marco A, Gonzalez Paz O, Fini I, Vignone D, Cellucci A, Battista M R, Auciello G, Orsatti L, Zini M, Monteagudo E, Khetarpal V, Rose M, Dominguez C, Herbst T, Toledo-Sherman L, Summa V, Munoz-Sanjuan I. (2019). Application of an in Vitro Blood-Brain Barrier Model in the Selection of Experimental Drug Candidates for the Treatment of Huntington’s Disease. Mol Pharm, 16(5), 2069–2082. 10.1021/acs.molpharmaceut.9b00042

Dong C, Ubogu E E. (2018). GDNF enhances human blood-nerve barrier function in vitro via MAPK signaling pathways. Tissue Barriers, 6(4), 1–22. 10.1080/21688370.2018.1546537

Eigenmann D E, Xue G, Kim K S, Moses A V, Hamburger M, Oufir M. (2013). Comparative study of four immortalized human brain capillary endothelial cell lines, hCMEC/D3, hBMEC, TY10, and BB19, and optimization of culture conditions, for an in vitro blood–brain barrier model for drug permeability studies. Fluids Barriers CNS, *10*(1), 33. 10.1186/2045-8118-10-33

Esaki T, Ohashi R, Watanabe R, Natsume-Kitatani Y, Kawashima H, Nagao C, Mizuguchi K. (2019). Computational Model To Predict the Fraction of Unbound Drug in the Brain. J Chem Inf Model, 59(7), 3251–3261. 10.1021/acs.jcim.9b00180

Fenstermacher J, Gross P, Sposito N, Acuff V, Pettersen S, Gruber K. (1988). Structural and Functional Variations in Capillary Systems within the Brain. Ann N Y Acad Sci, 529, 21–30. 10.1111/j.1749-6632.1988.tb51416.x

Fielder G C, Yang T W, Razdan M, Li Y, Lu J, Perry J K, Lobie P E, Liu D X. (2018). The GDNF Family: A Role in Cancer? Neoplasia, 20(1), 99–117. 10.1016/j.neo.2017.10.010

Friden M, Bergstrom F, Wan H, Rehngren M, Ahlin G, Hammarlund-Udenaes M, Bredberg U. (2011). Measurement of unbound drug exposure in brain: modeling of pH partitioning explains diverging results between the brain slice and brain homogenate methods. Drug Metab Dispos, 39(3), 353–362. 10.1124/dmd.110.035998

Fu J, Li L, Huo D, Zhi S, Yang R, Yang B, Xu B, Zhang T, Dai M, Tan C, Chen H, Wang X. (2021). Astrocyte-Derived TGFβ1 Facilitates Blood–Brain Barrier Function via Non-Canonical Hedgehog Signaling in Brain Microvascular Endothelial Cells. Brain Sciences, 11(1). 10.3390/brainsci11010077

Hanafy A S, Dietrich D, Fricker G, Lamprecht A. (2021). Blood-brain barrier models: Rationale for selection. Adv Drug Deliv Rev, 176, 113859. 10.1016/j.addr.2021.113859

Hashimoto Y, Greene C, Munnich A, Campbell M. (2023). The CLDN5 gene at the blood-brain barrier in health and disease. Fluids and Barriers of the CNS, 20(1). 10.1186/s12987-023-00424-5

Hatherell K, Couraud P O, Romero I A, Weksler B, Pilkington G J. (2011). Development of a three-dimensional, all-human in vitro model of the blood-brain barrier using mono-, co-, and tri-cultivation Transwell models. J Neurosci Methods, 199(2), 223–229. 10.1016/j.jneumeth.2011.05.012

He F, Wang Q F, Li L, Yu C, Liu C Z, Wei W C, Chen L P, Li H Y. (2023). Melatonin Protects Against Hyperoxia-Induced Apoptosis in Alveolar Epithelial type II Cells by Activating the MT2/PI3K/AKT/ETS1 Signaling Pathway. Lung, *201*(2), 225-234. 10.1007/s00408-023-00610-0

Hidalgo-Figueroa M, Bonilla S, Gutierrez F, Pascual A, Lopez-Barneo J. (2012). GDNF is predominantly expressed in the PV+ neostriatal interneuronal ensemble in normal mouse and after injury of the nigrostriatal pathway. J Neurosci, 32(3), 864–872. 10.1523/JNEUROSCI.2693-11.2012

Hui K, Wu S, Yue Y, Gu Y, Guan B, Wang X, Hsieh J T, Chang L S, He D, Wu K. (2018). RASAL2 inhibits tumor angiogenesis via p-AKT/ETS1 signaling in bladder cancer. Cell Signal, 48, 38–44. 10.1016/j.cellsig.2018.04.006

Igarashi Y, Utsumi H, Chiba H, Yamada-Sasamori Y, Tobioka H, Kamimura Y, Furuuchi K, Kokai Y, Nakagawa T, Mori M, Sawada N. (1999). Glial cell line-derived neurotrophic factor induces barrier function of endothelial cells forming the blood-brain barrier. Biochem Biophys Res Commun, 261(1), 108–112. 10.1006/bbrc.1999.0992

Ito R, Umehara K, Suzuki S, Kitamura K, Nunoya K I, Yamaura Y, Imawaka H, Izumi S, Wakayama N, Komori T, Anzai N, Akita H, Furihata T. (2019). A Human Immortalized Cell-Based Blood-Brain Barrier Triculture Model: Development and Characterization as a Promising Tool for Drug-Brain Permeability Studies. Mol Pharm, 16(11), 4461–4471. 10.1021/acs.molpharmaceut.9b00519

Ji-Ae K, RyojiI Y, Teruo N. (2009). IGF-1 released by corneal epithelial cells induces up-regulation of N-cadherin in corneal fibroblasts. J Cell Physiol, 221(1), 254–261. 10.1002/jcp.21850

Kodaira H, Kusuhara H, Fujita T, Ushiki J, Fuse E, Sugiyama Y. (2011). Quantitative evaluation of the impact of active efflux by p-glycoprotein and breast cancer resistance protein at the blood-brain barrier on the predictability of the unbound concentrations of drugs in the brain using cerebrospinal fluid concentration as a surrogate. J Pharmacol Exp Ther, 339(3), 935–944. 10.1124/jpet.111.180398

Lelièvre E, Mattot V, Huber P, Vandenbunde B, Soncin F. (2000). ETS1 lowers capillary endothelial cell density at confluence and induces the expression of VE-cadherin. Oncogene, 19(20), 2438–2446. 10.1038/sj.onc.1203563

Lelièvre E, Lionneton F, Soncin F, Vandenbunder B. (2001). The Ets family contains transcriptional activators and repressors involved in angiogenesis. Int J Biochem Cell Biol, 33(4), 391–407. 10.1016/s1357-2725(01)00025-5

Li P, Yang Y, Lin Z, Hong S, Jiang L, Zhou H, Yang L, Zhu L, Liu X, Liu L. (2022). Bile Duct Ligation Impairs Function and Expression of Mrp1 at Rat Blood-Retinal Barrier via Bilirubin-Induced P38 MAPK Pathway Activations. Int J Mol Sci, 23(14). 10.3390/ijms23147666

Li W, Chen Z, Chin I, Chen Z, Dai H. (2018). The Role of VE-cadherin in Blood-brain Barrier Integrity Under Central Nervous System Pathological Conditions. Current Neuropharmacology, 16(9), 1375–1384. 10.2174/1570159x16666180222164809

Li Y, Li Y, Zhang Y, Zhao Q, Zhang P, Sun M, Liu B, Yang H, Li P. (2022). Muscone and (+)-Borneol Cooperatively Strengthen CREB Induction of Claudin 5 in IL-1β-Induced Endothelium Injury. Antioxidants, 11(8). 10.3390/antiox11081455

Lin C H, Wang C H, Hsu S L, Liao L Y, Lin T A, Hsueh C M. (2016). Molecular Mechanisms Responsible for Neuron-Derived Conditioned Medium (NCM)-Mediated Protection of Ischemic Brain. PLoS One, 11(1), e0146692. 10.1371/journal.pone.0146692

Liu D, Yang L, Liu P, Ji X, Qi X, Wang Z, Chi T, Zou L. (2022). Sigma-1 receptor activation alleviates blood-brain barrier disruption post cerebral ischemia stroke by stimulating the GDNF-GFRalpha1-RET pathway. Exp Neurol, 347, 113867. 10.1016/j.expneurol.2021.113867

Liu X, Smith B J, Chen C, Callegari E, Becker S L, Chen X, Cianfrogna J, Doran A C, Doran S D, Gibbs J P, Hosea N, Liu J, Nelson F R, Szewc M A, Van Deusen J. (2005). Use of a physiologically based pharmacokinetic model to study the time to reach brain equilibrium: an experimental analysis of the role of blood-brain barrier permeability, plasma protein binding, and brain tissue binding. J Pharmacol Exp Ther, 313(3), 1254–1262. 10.1124/jpet.104.079319

Liu Y, Mu S, Li X, Liang Y, Wang L, Ma X. (2019). Unfractionated Heparin Alleviates Sepsis-Induced Acute Lung Injury by Protecting Tight Junctions. Journal of Surgical Research, 238, 175–185. 10.1016/j.jss.2019.01.020

Lonka-Nevalaita L, Lume M, Leppänen S, Jokitalo E, Peränen J, Saarma M. (2010). Characterization of the Intracellular Localization, Processing, and Secretion of Two Glial Cell Line-Derived Neurotrophic Factor Splice Isoforms. The Journal of Neuroscience, 30(34), 11403–11413. 10.1523/jneurosci.5888-09.2010

Luo Y, Yang H, Wan Y, Yang S, Wu J, Chen S, Li Y, Jin H, He Q, Zhu D Y, Zhou Y, Hu B. (2022). Endothelial ETS1 inhibition exacerbate blood-brain barrier dysfunction in multiple sclerosis through inducing endothelial-to-mesenchymal transition. Cell Death Dis, 13(5), 462. 10.1038/s41419-022-04888-5

Mathiisen T M, Lehre K P, Danbolt N C, Ottersen O P. (2010). The perivascular astroglial sheath provides a complete covering of the brain microvessels: an electron microscopic 3D reconstruction. Glia, 58(9), 1094–1103. 10.1002/glia.20990

Maurer T S, Debartolo D B, Tess D A, Scott D O. (2005). Relationship between exposure and nonspecific binding of thirty-three central nervous system drugs in mice. Drug Metab Dispos, 33(1), 175–181. 10.1124/dmd.104.001222

Ming-Shan L, Juan C, Ling H, Lu L, Bian-Sheng J. (2013). CJY, an isoflavone, interacts with ATPase of P-glycoprotein in the rat brain microvessel endothelial cells (RBMECs). J Chemother, 25(6), 347–354. 10.1179/1973947813Y.0000000094

Morita A, Yamashita N, Sasaki Y, Uchida Y, Nakajima O, Nakamura F, Yagi T, Taniguchi M, Usui H, Katoh-Semba R, Takei K, Goshima Y. (2006). Regulation of Dendritic Branching and Spine Maturation by Semaphorin3A-Fyn Signaling. The Journal of Neuroscience, 26(11), 2971–2980. 10.1523/jneurosci.5453-05.2006

Muoio V, Persson P B, Sendeski M M. (2014). The neurovascular unit - concept review. Acta Physiol, 210(4), 790–798. 10.1111/apha.12250

Nakagawa S, Deli M A, Kawaguchi H, Shimizudani T, Shimono T, Kittel A, Tanaka K, Niwa M. (2009). A new blood-brain barrier model using primary rat brain endothelial cells, pericytes and astrocytes. Neurochem Int, 54(3-4), 253–263. 10.1016/j.neuint.2008.12.002

Nitta T, Hata M, Gotoh S, Seo Y, Sasaki H, Hashimoto N, Furuse M, Tsukita S. (2003). Size-selective loosening of the blood-brain barrier in claudin-5-deficient mice. J Cell Biol, 161(3), 653–660. 10.1083/jcb.200302070

Nowrangi D S, McBride D, Manaenko A, Dixon B, Tang J, Zhang J H. (2019). rhIGF-1 reduces the permeability of the blood-brain barrier following intracerebral hemorrhage in mice. Exp Neurol, 312, 72–81. 10.1016/j.expneurol.2018.11.009

Ohtsuki S, Sato S, Yamaguchi H, Kamoi M, Asashima T, Terasaki T. (2007). Exogenous expression of claudin-5 induces barrier properties in cultured rat brain capillary endothelial cells. J Cell Physiol, 210(1), 81–86. 10.1002/jcp.20823

Ohtsuki S, Ikeda C, Uchida Y, Sakamoto Y, Miller F, Glacial F, Decleves X, Scherrmann J M, Couraud P O, Kubo Y, Tachikawa M, Terasaki T. (2013). Quantitative targeted absolute proteomic analysis of transporters, receptors and junction proteins for validation of human cerebral microvascular endothelial cell line hCMEC/D3 as a human blood-brain barrier model. Mol Pharm, 10(1), 289–296. 10.1021/mp3004308

Palmiotti C A, Prasad S, Naik P, Abul K M, Sajja R K, Achyuta A H, Cucullo L. (2014). In vitro cerebrovascular modeling in the 21st century: current and prospective technologies. Pharm Res, 31(12), 3229–3250. 10.1007/s11095-014-1464-6

Pochon N A, Menoud A, Tseng J L, Zurn A D, Aebischer P. (1997). Neuronal GDNF Expression in the Adult Rat Nervous System Identified By In Sifu Hybridization. European Journal of Neuroscience, 9(3), 463–471. 10.1111/j.1460-9568.1997.tb01623.x

Potjewyd G, Moxon S, Wang T, Domingos M, Hooper N M. (2018). Tissue Engineering 3D Neurovascular Units: A Biomaterials and Bioprinting Perspective. Trends Biotechnol, 36(4), 457–472. 10.1016/j.tibtech.2018.01.003

Qi D, Lin H, Hu B, Wei Y. (2023). A review on in vitro model of the blood-brain barrier (BBB) based on hCMEC/D3 cells. J Control Release, 358, 78–97. 10.1016/j.jconrel.2023.04.020

Sajja R K, Prasad S, Cucullo L. (2014). Impact of altered glycaemia on blood-brain barrier endothelium: an in vitro study using the hCMEC/D3 cell line. Fluids Barriers CNS, 11(1), 8. 10.1186/2045-8118-11-8

Saker S, Stewart E A, Browning A C, Allen C L, Amoaku W M. (2014). The effect of hyperglycaemia on permeability and the expression of junctional complex molecules in human retinal and choroidal endothelial cells. Exp Eye Res, 121, 161–167. 10.1016/j.exer.2014.02.016

Sanchez-Dengra B, Gonzalez-Alvarez I, Sousa F, Bermejo M, Gonzalez-Alvarez M, Sarmento B. (2021). In vitro model for predicting the access and distribution of drugs in the brain using hCMEC/D3 cells. Eur J Pharm Biopharm, 163, 120–126. 10.1016/j.ejpb.2021.04.002

Sauteur L, Affolter M, Belting H-G. (2017). Distinct and redundant functions of Esam and VE-cadherin during vascular morphogenesis. Development. 10.1242/dev.140038

Savettieri G, Di Liegro I, Catania C, Licata L, Pitarresi G L, D’Agostino S, Gabriella S, De Caro V, Giandalia G, Giannola L I, Cestelli A. (2000). Neurons and ECM regulate occludin localization in brain endothelial cells. Neuroreport, 11(5), 1081–1084. 10.1097/00001756-200004070-00035

Schiera G, Bono E, Raffa M P, Gallo A, Pitarresi G L, Di Liegro I, Savettieri G. (2003). Synergistic effects of neurons and astrocytes on the differentiation of brain capillary endothelial cells in culture. J Cell Mol Med, 7(2), 165–170. 10.1111/j.1582-4934.2003.tb00215.x

Schiera G, Sala S, Gallo A, Raffa M P, Pitarresi G L, Savettieri G, Di Liegro I. (2005). Permeability properties of a three-cell type in vitro model of blood-brain barrier. Journal of Cellular and Molecular Medicine, 9(2), 373–379. 10.1111/j.1582-4934.2005.tb00362.x

Shimizu F, Sano Y, Abe M A, Maeda T, Ohtsuki S, Terasaki T, Kanda T. (2011). Peripheral nerve pericytes modify the blood-nerve barrier function and tight junctional molecules through the secretion of various soluble factors. J Cell Physiol, 226(1), 255–266. 10.1002/jcp.22337

Shimizu F, Sano Y, Saito K, Abe M A, Maeda T, Haruki H, Kanda T. (2012). Pericyte-derived glial cell line-derived neurotrophic factor increase the expression of claudin-5 in the blood-brain barrier and the blood-nerve barrier. Neurochem Res, 37(2), 401–409. 10.1007/s11064-011-0626-8

Summerfield S G, Read K, Begley D J, Obradovic T, Hidalgo I J, Coggon S, Lewis A V, Porter R A, Jeffrey P. (2007). Central nervous system drug disposition: the relationship between in situ brain permeability and brain free fraction. J Pharmacol Exp Ther, 322(1), 205–213. 10.1124/jpet.107.121525

Sweeney M D, Zhao Z, Montagne A, Nelson A R, Zlokovic B V. (2019). Blood-Brain Barrier: From Physiology to Disease and Back. Physiol Rev, 99(1), 21–78. 10.1152/physrev.00050.2017

Taddei A, Giampietro C, Conti A, Orsenigo F, Breviario F, Pirazzoli V, Potente M, Daly C, Dimmeler S, Dejana E. (2008). Endothelial adherens junctions control tight junctions by VE-cadherin-mediated upregulation of claudin-5. Nat Cell Biol, 10(8), 923–934. 10.1038/ncb1752

Tang E D, Nuñez G, Barr F G, Guan K-L. (1999). Negative Regulation of the Forkhead Transcription Factor FKHR by Akt. Journal of Biological Chemistry, 274(24), 16741–16746. 10.1074/jbc.274.24.16741

Tavelin S, Gråsjö J, Taipalensuu J, Ocklind G, Artursson P. (2002). Applications of Epithelial Cell Culture in Studies of Drug Transport. In C. Wise (Ed.), Epithelial Cell Culture Protocols (pp. 233-272). Totowa, NJ: Humana Press. 10.1385/1-59259-185-x:233

Tunggal J A, Helfrich I, Schmitz A, Schwarz H, Günzel D, Fromm M, Kemler R, Krieg T, Niessen C M. (2005). E-cadherin is essential for in vivo epidermal barrier function by regulating tight junctions. The EMBO Journal, 24(6), 1146–1156. 10.1038/sj.emboj.7600605

Wang Z G, Cheng Y, Yu X C, Ye L B, Xia Q H, Johnson N R, Wei X, Chen D Q, Cao G, Fu X B, Li X K, Zhang H Y, Xiao J. (2016). bFGF Protects Against Blood-Brain Barrier Damage Through Junction Protein Regulation via PI3K-Akt-Rac1 Pathway Following Traumatic Brain Injury. Mol Neurobiol, 53(10), 7298–7311. 10.1007/s12035-015-9583-6

Watanabe D, Takagi H, Suzuma K, Suzuma I, Oh H, Ohashi H, Kemmochi S, Uemura A, Ojima T, Suganami E, Miyamoto N, Sato Y, Honda Y. (2004). Transcription factor Ets-1 mediates ischemia- and vascular endothelial growth factor-dependent retinal neovascularization. Am J Pathol, 164(5), 1827–1835. 10.1016/S0002-9440(10)63741-8

Watanabe D, Nakagawa S, Morofuji Y, Toth A E, Vastag M, Aruga J, Niwa M, Deli M A. (2021). Characterization of a Primate Blood-Brain Barrier Co-Culture Model Prepared from Primary Brain Endothelial Cells, Pericytes and Astrocytes. Pharmaceutics, 13(9). 10.3390/pharmaceutics13091484

Weber M L, Hofland C M, Shaffer C L, Flik G, Cremers T, Hurst R S, Rollema H. (2013). Therapeutic doses of antidepressants are projected not to inhibit human alpha4beta2 nicotinic acetylcholine receptors. Neuropharmacology, 72, 88–95. 10.1016/j.neuropharm.2013.04.027

Weksler B, Romero I A, Couraud P-O. (2013). The hCMEC/D3 cell line as a model of the human blood brain barrier. Fluids and Barriers of the CNS, 10(1), 16. 10.1186/2045-8118-10-16

Wu T, Sheng Y, Qin Y Y, Kong W M, Jin M M, Yang H Y, Zheng X K, Dai C, Liu M, Liu X D, Liu L. (2021). Bile duct ligation causes opposite impacts on the expression and function of BCRP and P-gp in rat brain partly via affecting membrane expression of ezrin/radixin/moesin proteins. Acta Pharmacol Sin, 42(11), 1942–1950. 10.1038/s41401-020-00602-3

Xue Q, Liu Y, Qi H, Ma Q, Xu L, Chen W, Chen G, Xu X. (2013). A Novel Brain Neurovascular Unit Model with Neurons, Astrocytes and Microvascular Endothelial Cells of Rat. International Journal of Biological Sciences, 9(2), 174–189. 10.7150/ijbs.5115

Yang H, Su M, Liu M, Sheng Y, Zhu L, Yang L, Mu R, Zou J, Liu X, Liu L. (2023). Hepatic retinaldehyde deficiency is involved in diabetes deterioration by enhancing PCK1- and G6PC-mediated gluconeogenesis. Acta Pharm Sin B, 13(9), 3728–3743. 10.1016/j.apsb.2023.06.014

Zhang X, Tang N, Hadden T J, Rishi A K. (2011). Akt, FoxO and regulation of apoptosis. Biochimica et Biophysica Acta, 1813(11), 1978–1986. 10.1016/j.bbamcr.2011.03.010

Zhou L, Schmidt K, Nelson F R, Zelesky V, Troutman M D, Feng B. (2009). The effect of breast cancer resistance protein and P-glycoprotein on the brain penetration of flavopiridol, imatinib mesylate (Gleevec), prazosin, and 2-methoxy-3-(4-(2-(5-methyl-2-phenyloxazol-4-yl)ethoxy)phenyl)propanoic acid (PF-407288) in mice. Drug Metab Dispos, 37(5), 946–955. 10.1124/dmd.108.024489

Zhou Y, Zhou J, Li P, Xie Q, Sun B, Li Y, Chen Y, Zhao K, Yang T, Zhu L, Xu J, Liu X, Liu L. (2019). Increase in P-glycoprotein levels in the blood-brain barrier of partial portal vein ligation /chronic hyperammonemia rats is medicated by ammonia/reactive oxygen species/ERK1/2 activation: In vitro and in vivo studies. Eur J Pharmacol, 846, 119–127. 10.1016/j.ejphar.2019.01.005

